# Ultrasound system for precise neuromodulation of human deep brain circuits

**DOI:** 10.1101/2024.06.08.597305

**Authors:** Eleanor Martin, Morgan Roberts, Ioana F Grigoras, Olivia Wright, Tulika Nandi, Sebastian W Rieger, Jon Campbell, Tim den Boer, Ben T Cox, Charlotte J Stagg, Bradley E Treeby

## Abstract

Transcranial ultrasound stimulation (TUS) has emerged as a promising technique for non-invasive neuromodulation, but current systems lack the precision to target deep brain structures effectively. Here, we introduce an advanced TUS system that achieves unprecedented precision in deep brain neuromodulation. The system features a 256-element, helmet-shaped transducer array operating at 555 kHz, coupled with a stereotactic positioning system, individualised treatment planning, and real-time monitoring using functional MRI. In a series of experiments, we demonstrate the system’s ability to selectively modulate the activity of the lateral geniculate nucleus (LGN) and its functionally connected regions in the visual cortex. Participants exhibited significantly increased visual cortex activity during concurrent TUS and visual stimulation, with high reproducibility across individuals. Moreover, a theta-burst TUS protocol induced robust neuromodulatory effects, with decreased visual cortex activity observed for at least 40 minutes post-stimulation. These neuromodulatory effects were specific to the targeted LGN, as confirmed by control experiments. Our findings highlight the potential of this advanced TUS system to non-invasively modulate deep brain circuits with high precision and specificity, offering new avenues for studying brain function and developing targeted therapies for neurological and psychiatric disorders. The unprecedented spatial resolution and prolonged neuromodulatory effects demonstrate the transformative potential of this technology for both research and clinical applications, paving the way for a new era of non-invasive deep brain neuromodulation.

---

Deep within the human brain are a group of grey matter structures, the basal ganglia and thalamic nuclei, which play pivotal roles in all aspects of human behaviour. Indeed, their dysregulation is pathognomonic of numerous neurological and psychiatric conditions.^1^ The ability to precisely modulate neuronal activity within these areas offers potentially revolutionary therapeutic avenues for these often devastating disorders that are resistant to traditional treatments.^2^ Furthermore, it unlocks insights into neural circuitry in healthy brains, presenting a plausible route to breakthrough shifts in our understanding of fundamental cognitive processes such as consciousness.^3^

However, current neuromodulation techniques face significant limitations in targeting deep brain structures. Deep Brain Stimulation (DBS), though effective, is invasive and carries surgical risks.^4^ Transcranial Magnetic Stimulation (TMS) and Transcranial Direct Current Stimulation (tDCS) offer non-invasive alternatives but lack the requisite depth penetration and spatial precision. TMS primarily influences cortical areas, and while its variant, Deep TMS, attempts deeper reach, it still falls short of the precision needed for specific deep brain targets.^5^ tDCS, and the related technique of temporal interference, are even less focused and more diffuse, making targeted deep brain modulation a challenge.^6^

Transcranial Ultrasound Stimulation (TUS) has emerged as a promising modality for non-invasive brain modulation, offering the unique advantage of deep tissue penetration.^7,8^ This technique, leveraging the application of ultrasound waves, holds substantial potential for influencing neural activity in humans in both superficial and deep brain regions. However, a significant limitation of existing TUS systems, which typically employ small aperture transducers, is the compromise in focal precision. While these systems can reach deep brain structures, the spatial resolution of the stimulation is often suboptimal, potentially affecting a broader region than intended.^9,10^ This inherent trade-off between depth penetration and focal size underscores the need for advanced TUS systems capable of delivering more localised and precise neuromodulation in humans, particularly in the context of targeting deep brain structures such as the thalamus.

In addressing the limitations of focal precision in TUS, large hemispherical arrays have shown promise, particularly as evidenced in MR-guided Focused Ultrasound (MRgFUS). These arrays contain numerous transducer elements over a large aperture allowing for finer control over the ultrasound beam with a significantly reduced focal size.^11^ This technology has been successfully applied for ablative therapies, such as targeting the ventral intermediate nucleus (VIM) of the thalamus for treating essential tremor.^12^ Recently, a study showed that low-power, non-thermal stimulation of the VIM and the dentato-rubro-thalamic tract (DRT) using an MRgFUS array could induce a sustained reduction of essential tremor in patients.^13^ However, these arrays rely on positioning the patient using a neurosurgical frame, positioned using skull screws, and focal tissue heating for target confirmation, making them unsuitable for non-invasive, reversible neuromodulation in healthy individuals. To date, no system has been available for neuroscientific studies that can non-invasively modulate activity in the deep brain with the spatial precision required to target individual thalamic nuclei.

Here, we introduce an advanced transcranial ultrasound system that achieves unprecedented precision in human deep brain neuromodulation. The system features a 256-element sparse array within an ellipsoidal helmet, enabling focal stimulation in deep brain areas. Uniquely, it is compatible with simultaneous fMRI imaging, allowing for real-time monitoring of neuromodulatory effects. A custom-designed stereotactic face and neck mask ensures precise participant positioning, while a model-based treatment planning method and an online re-planning mechanism maintain accurate targeting. This eliminates the need for a surgical frame for participant positioning and focal tissue heating for target confirmation, for the first time making a high precision deep brain neuromodulation system available for study of the healthy brain.

We demonstrate the system’s efficacy through two rigorously-designed experiments in healthy human participants, targeting the lateral geniculate nucleus (LGN), one of the smallest functionally distinct nuclei of the thalamus. These experiments reveal significant and specific network effects of TUS within connected brain regions, as evidenced by changes in network activity measured using fMRI during and after task performance. Our findings underscore the potential of this advanced transcranial ultrasound system to revolutionise deep brain neuromodulation, offering new avenues for studying brain function and treating neurological and psychiatric disorders.

## Ultrasound system for precise modulation of deep brain structures

We developed an advanced transcranial ultrasound system designed for highly focal modulation of deep brain structures inside an MR scanner (Figure 1, Extended Data Figure 2). The system is based around a semi-ellipsoidal helmet housing 256 individually-controllable transducer elements operating at a frequency of 555 kHz. A water coupling system with temperature control and hydrostatic pressure compensation ensures efficient energy transfer to the head. The helmet’s dimensions and angle were optimised based on an analysis of the average adult head size to ensure comfort, accommodate a wide range of head sizes, and minimise the distance and angle of incidence to the head (Extended Data Figure 1). Numerical simulations were employed to determine the optimal element configuration within the helmet, balancing focal size and grating lobe levels while maintaining line of sight between the element positions and deep brain structures.

**Figure 1:**
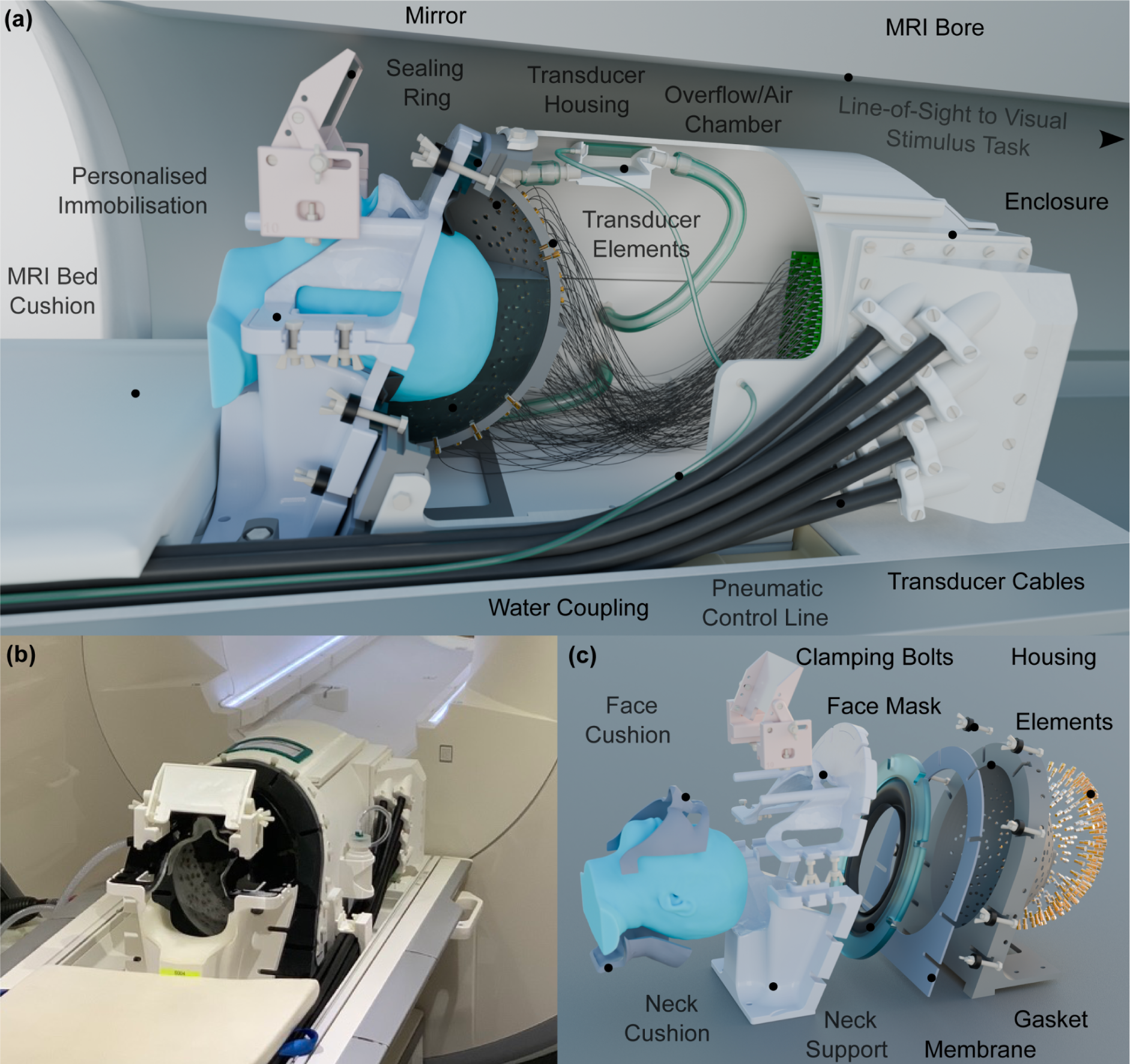
**(a)** Advanced transcranial ultrasound system within the MR bore, used for concurrent neuromodulation and functional neuroimaging. The participant is immobilised and coupled to the transducer array using water. The participant has line of sight to a visual stimulus task outside of the bore, via a mirror. **(b)** System positioned on the MR table. **(c)** Exploded view of the personalised immobilisation hardware, showing the sealing membrane and gaskets which retain water between the participant’s head and transducer bowl.

To characterise the system’s performance, we conducted comprehensive acoustic measurements and simulations (Extended Data Figure 3). The system demonstrated the ability to steer over a wide range centred on the helmet’s geometric centre, with a −3 dB focal size of 1.3 mm laterally and 3.4 mm axially at the geometric focus giving a focal volume of 24 mm^3^ which is maintained across an extremely wide range of target locations (Figure 2b, Extended Data Figure 3). Notably, this focal size is approximately 1000 times smaller than that achieved by conventional small aperture ultrasound transducers,^9,14^ and 30 times smaller than devices previously designed specifically for deep brain targeting in healthy humans.^15^ Comprehensive compatibility testing demonstrated negligible impact of the ultrasound system on MR image quality and no influence of the MR environment on acoustic output (Extended Data Figure 4). A synchronisation setup was implemented to interleave ultrasound and MR acquisitions, effectively mitigating electromagnetic interference during simultaneous operation.

**Figure 2:**
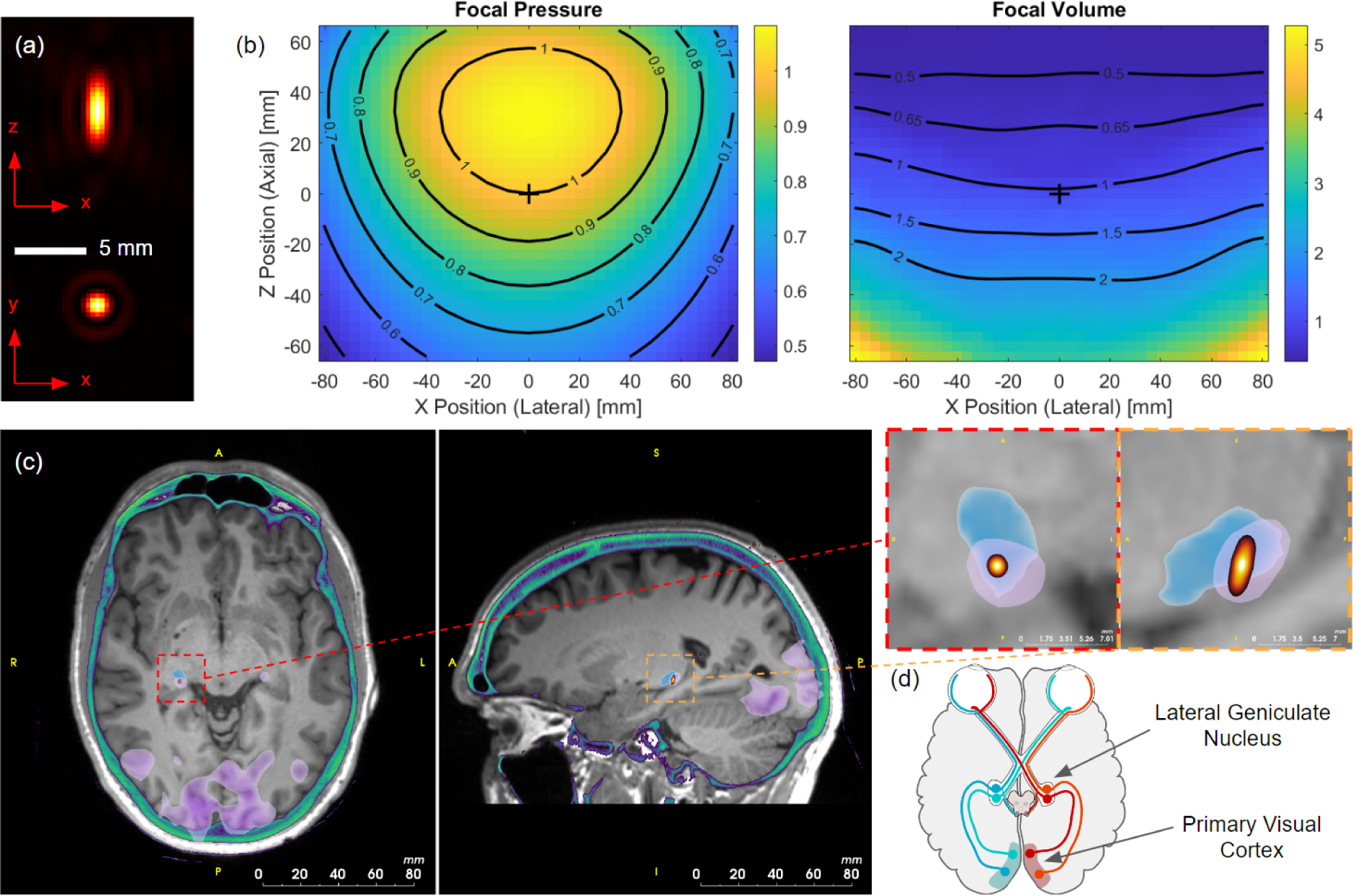
**(a)** Focal intensity for the transcranial ultrasound system in the axial (top) and lateral (bottom) planes at the geometric focus. **(b)** Relative focal pressure and −3dB focal volume as a function of lateral and axial steering position. The values are normalised to the focal pressure and focal volume at the geometric centre of the helmet (position shown with a + symbol). The highly targeted focus is maintained over a very wide steering range. **(c)** Planning images showing the T1-weighted MR (grayscale), CT image thresholded to show the skull (green), functional activation from visual task (purple), statistical segmentation of the lateral geniculate nucleus (LGN) (blue), and −6 dB volume of the simulated target pressure (yellow/red). **(d)** Schematic of the visual system showing the connections between the left and right LGN and the primary visual cortices (adapted from ’Human Visual Pathway’, retrieved from https://commons.wikimedia.org/wiki/File:Human_visual_pathway.svg).

The system’s small focal spot (Figure 2) enables high-precision targeting of deep brain structures but necessitates precise alignment between the participant’s head and the transducer array to ensure accurate stimulation of the desired brain region. To address this challenge, we developed a custom-designed stereotactic face and neck mask derived from individual participant MR data that can comfortably be used for healthy participants (Figure 1, Extended Data Figure 2). The mask, fabricated using 3D printing and casting techniques, comprises two parts: a neck support and a face mask, each designed to engage specific anatomical landmarks while leaving openings for the participant’s eyes, ears, and mouth. The mask is securely attached to the helmet using quick-release connectors, ensuring consistent alignment between the participant and the transducer array. This approach demonstrated high inter-session positioning repeatability, with a mean target shift of 1.50 +/-0.70 mm across participants and sessions, and high intra-session stability, with an average participant motion of 0.25 +/-0.001 mm during scans. An adjustable mirror system was also integrated into the mask, enabling participants to view a visual display unit positioned at the end of the MRI bore during experiments.

Beyond positioning, precise targeting of deep brain structures requires a treatment planning approach that accounts for the aberration and attenuation caused by the skull. We employed k-Plan, our commercially available software, to prospectively compute the driving parameters for each transducer element based on a full-wave acoustic model incorporating participant-specific skull and brain properties derived from low-dose CT scans (Figure 2c). To maintain targeting accuracy across sessions despite small shifts in participant position, we implemented an on-line re-planning protocol. This protocol adjusted the pre-calculated driving phases for each element by applying geometrically calculated offsets, effectively shifting the focus to align with the desired target. Experimental validation using human skull caps demonstrated that the measured focal parameters were within 21% for target pressure, 0.9 mm for position, and (dx, dy, dz) = (0.2, 0.2, 0.7) mm for the −3 dB focal dimensions compared to the treatment plan (Extended Data Figure 6f). Re-planning validation showed even better agreement, confirming the validity of the isoplanatic assumption for small positional adjustments (Extended Data Figure 6g).

## Significant target engagement and prolonged network effects with TUS

To demonstrate the capabilities of our advanced transcranial ultrasound system for precise neuromodulation of deep brain structures we targeted the well-characterised visual system. This network is an ideal testbed, as it involves both the lateral geniculate nucleus (LGN), a small, deep-brain structure, and the primary visual cortex (V1), a larger, cortical region that is monosynaptically connected to the LGN (see Figure 2d). With a volume of approximately 80 mm³, the LGN is well-suited to showcase the precise targeting capabilities of our ultrasound system, which has a −3 dB focal volume of 24 mm³ (see Figure 2c). Simultaneously, activity in V1 is readily observable using functional magnetic resonance imaging (fMRI), providing a reliable readout of network-level effects. We employed a visual checkerboard task to activate the visual system, minimising confounds associated with participant motion or additional equipment requirements. This experimental design allowed us to demonstrate the system’s ability to precisely modulate deep brain activity and observe its consequences on connected cortical regions.

We conducted two experiments on seven healthy participants using a dense-sampling approach, which involves comprehensive assessments within participants to gain detailed insights into the effects of TUS. This approach was chosen to provide a rigorous understanding of both immediate (online) and lasting (offline) neuromodulatory effects, aligning with the two main types of ultrasound experiments performed in previous TUS literature.

In the first experiment, we wanted to demonstrate target engagement with TUS using an online paradigm. We, therefore, hypothesised that active TUS to the LGN would lead to significant modulation of visually-related activity in the primary visual cortex. To test this hypothesis, we employed a single-blind, pseudo-randomized, sham-controlled block design. Participants fixated on a central point while a visual checkerboard stimulus was presented on a screen. Active TUS (300 ms pulses every 3 seconds, target pressure of 775 kPa; see Extended Data Figure 7) was applied during half of the visual stimulation blocks, while the other half served as sham stimulation, with the order of blocks pseudo-randomised to prevent order effects. Functional MRI volumes were acquired every 3 seconds, interleaved with the TUS pulses. Seven participants participated in two stimulation days, with six on-target stimulation runs per day. Additionally, three off-target stimulation runs were conducted for each of the first three participants, with the TUS focus targeted at the medial dorsal nucleus (MDN), a control site close to the LGN. This rigorous experimental design allowed for a robust comparison of the effects of active TUS versus sham stimulation on both the targeted LGN and its functionally connected V1.

During online stimulation, we demonstrated significantly increased task-related activity in the ipsilateral primary visual cortex during active TUS compared with sham across all participants (Figure 3). This finding provides compelling evidence for the effective target engagement of the LGN using our advanced transcranial ultrasound system. Importantly, there were no regions of significantly altered task-related activity during LGN stimulation within the contralateral visual cortex of the task-activated network, underlining the anatomical precision of our approach and the ability to selectively modulate the targeted deep brain structure without off-target effects.

**Figure 3:**
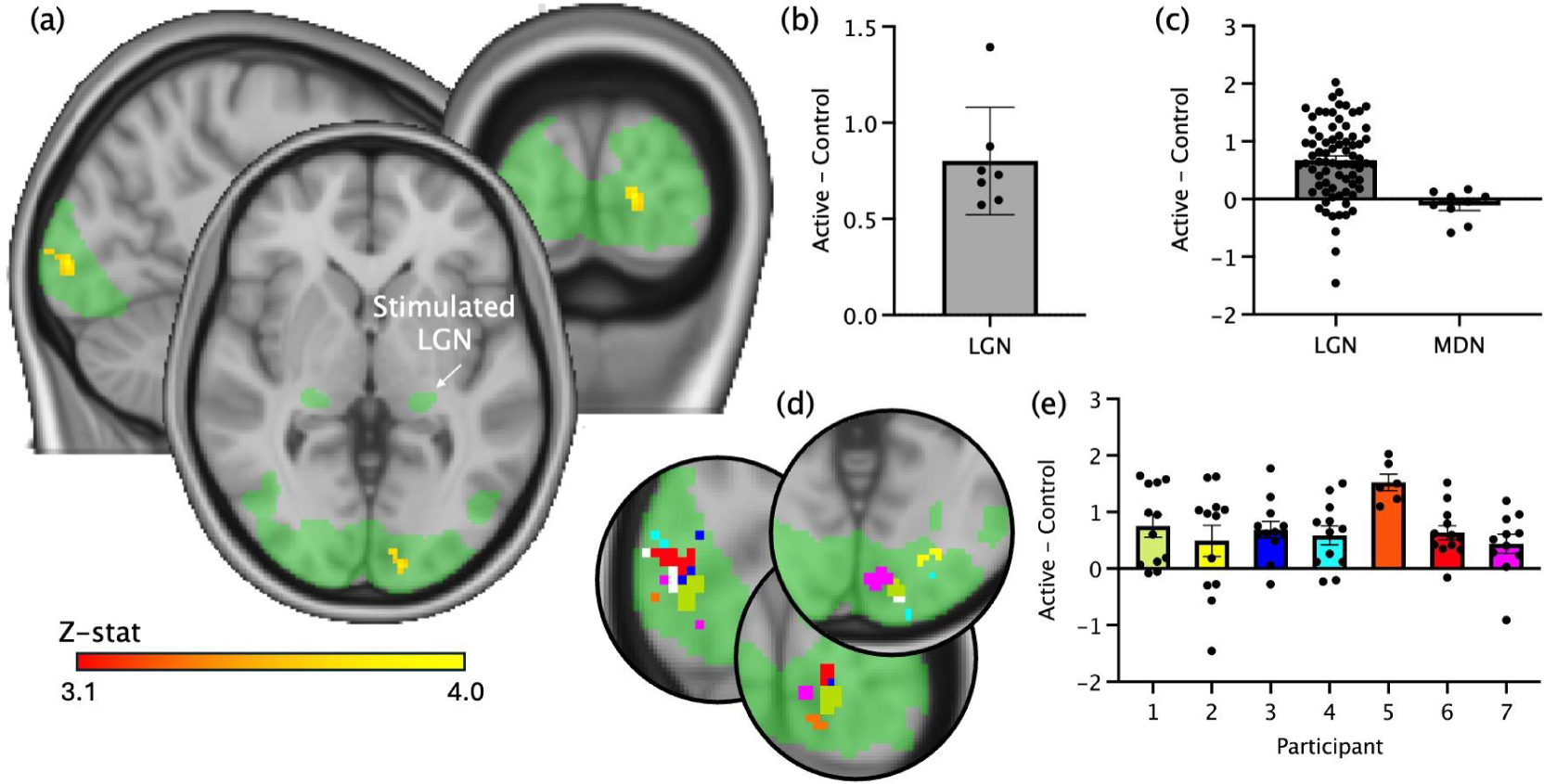
Ultrasound stimulation to the left LGN during a visual task leads to significantly increased activity in the ipsilateral occipital cortex compared with sham. **(a)** Group mean visually-related activation (green) is evident in the bilateral LGN and associated primary visual cortices during the task. Stimulation of the left (highlighted) LGN results in significantly increased task-related activity in a spatially-precise region within the directly connected ipsilateral visual cortex. No changes in activity are observed in the contralateral visual cortex (Mixed-effects, cluster-corrected with a threshold of z=3.1, p<0.05, red-yellow). **(b)** Difference in task-related activity (difference in z-scored BOLD task-related change) during LGN stimulation compared with sham within the peak task-related activation region in (a) for each participant. **(c)** Difference in task-related activity (difference in z-scored BOLD task-related change) during LGN stimulation (left) and MDN stimulation (right) compared with sham within the peak task-related activation region for each run separately. **(d)** The peak change in task-related activity during LGN ultrasound stimulation was highly similar across individual participants. Green: mean task-related activation, white: group mean results (as in (a)). **(e)** Difference in task-related activity (difference in z-scored BOLD task-related change) during LGN stimulation for each run, for each participant separately. Participant 5 only completed 6 runs and participant 7 11 runs.

To further validate the specificity of the observed effects, for three of the participants we conducted control experiments targeting the MDN, a thalamic nucleus close to LGN, as an active control site. TUS to this control site resulted in no significant changes in the visually-related activity within the ipsilateral occipital cortex, either on a whole-brain analysis or within the region significantly modulated by LGN stimulation, confirming that the modulation of V1 activity was indeed a consequence of precise LGN targeting and not a non-specific effect of ultrasound stimulation (Figure 3c). The results of LGN stimulation were remarkably similar across our seven participants, both in terms of anatomical location and effect size (Figure 3d-e). These results demonstrate the power of our advanced transcranial ultrasound system to precisely modulate deep brain activity and its potential to elucidate the functional roles of specific neural circuits in the human brain.

In the second experiment, we aimed to establish whether TUS to the LGN could lead to long-lasting after-effects in network activity, building upon the findings of the online stimulation protocol. We employed a within-participant design in four participants with an active control site to investigate this, applying TUS using a theta-patterned approach, which has previously been shown to induce prolonged changes in cortical excitability.^16–19^ This stimulation protocol, known as theta burst stimulation (TBS), consists of short, 20 ms ultrasound pulses delivered at a theta rhythm (5 Hz) for a total of 80 s. To measure brain response, participants underwent several fMRI scans during the visual checkerboard task: a baseline measurement before stimulation, an early post-stimulation task scan (scanning started 19-21 minutes after stimulation and lasted 16 minutes), and a late post-stimulation scans (scanning started approximately 140 minutes after stimulation and lasted 16 minutes). We hypothesised that TBS applied to the LGN would result in sustained modulation of visually-evoked activity in the ipsilateral primary visual cortex.

Our results revealed that active TUS to the LGN led to a significant decrease in visually-evoked activity in the ipsilateral primary visual cortex 40 minutes after stimulation compared to baseline (Figure 4). This change in activity was no longer significant at the later post-stimulation timepoint. The anatomical location of this significantly decreased activity was very similar to the region of significant response to the online TUS, further supporting the spatial specificity of the targeted stimulation. No significant changes in brain activity in the primary visual cortex were observed after offline stimulation of the active control site compared to baseline either at the early or late post-stimulation timepoints. These findings demonstrate the potential of our advanced transcranial ultrasound system to induce lasting plasticity in deep brain structures and their associated networks, opening new avenues for studying the mechanisms of neural adaptation and for developing novel therapeutic interventions.

**Figure 4:**
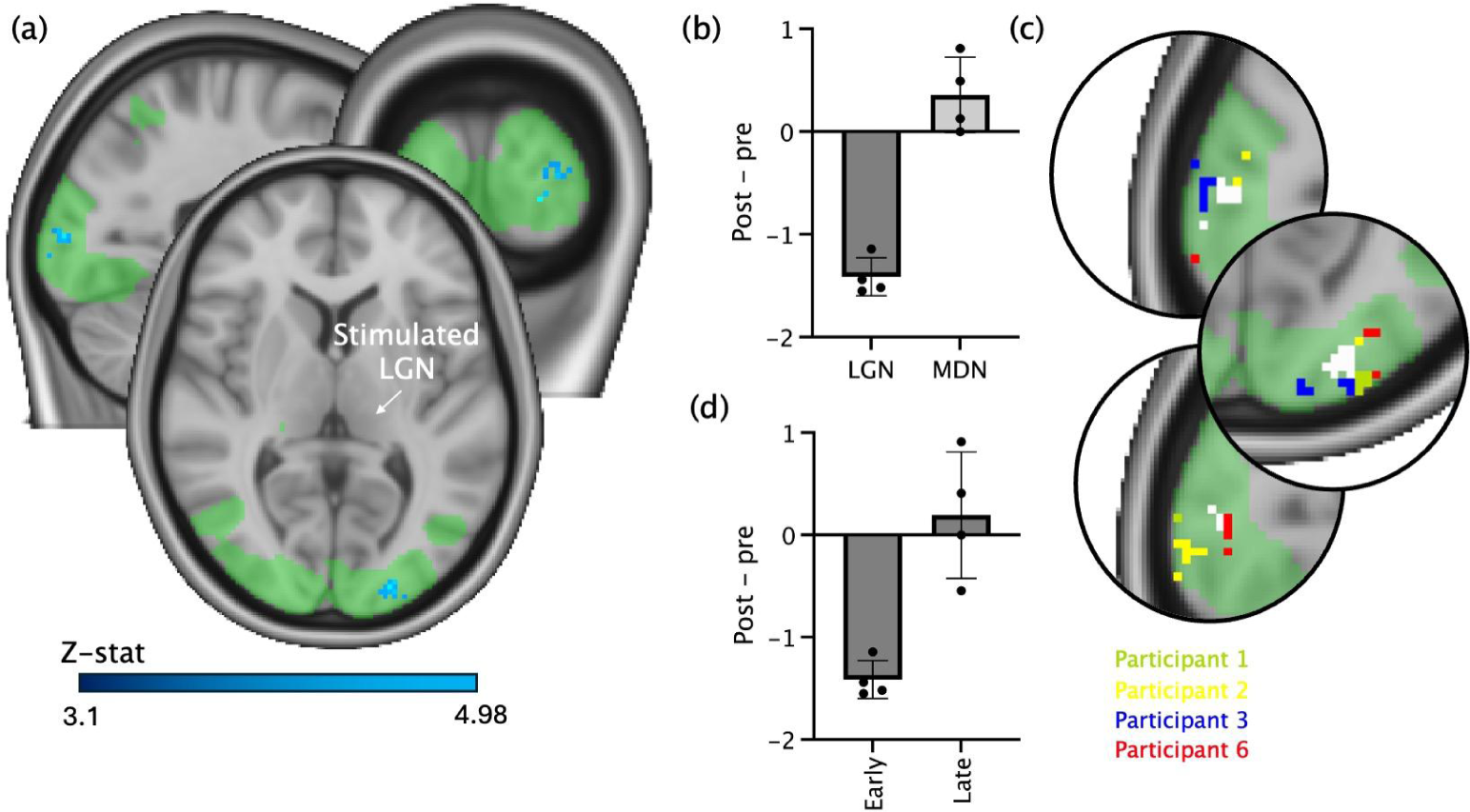
Offline ultrasound stimulation to the left LGN significantly decreases task-related activity in the ipsilateral visual cortex during a visual checkerboard task early after stimulation. **(a)** Stimulation of the left (highlighted) LGN results in significantly decreased task-related activity in the directly connected ipsilateral visual cortex but not in the contralateral visual cortex (blue-light blue; mixed effects, z=3.1, p< 0.05). These significant areas overlap the mean cortical activation map of the occipital cortex during all task blocks (green). **(b)** Change in task-related activity (difference in z-scored BOLD task-related change) after LGN (left) and MDN (right) stimulation within the peak task-related activation region in (a) for each run separately. **(c)** The peak change in task-related activity during LGN ultrasound stimulation was highly similar across individual participants. Green: mean task-related activation, white: group mean results (as in (a)). **(d)** Change in task-related activity (difference in z-scored BOLD task-related change) after LGN stimulation within the peak task-related activation region in (a) for each run separately, at the early and late time points.

## Discussion

In this study, we introduce a highly advanced transcranial ultrasound system that achieves unprecedented precision for deep brain neuromodulation. For the first time in humans, we demonstrate that this technology allows for specific targeting of individual thalamic nuclei non-invasively, paving the way for a new era in human neuroscience research. Through online stimulation, we provide evidence of target engagement by demonstrating that TUS of the LGN leads to a significant increase in activity in the downstream primary visual cortex. Moreover, using a stimulation paradigm designed to induce after-effects, we observe significant modulation of activity in the ipsilateral primary visual cortex for at least 40 minutes following the stimulation period. These findings represent a major step forward in our ability to precisely modulate deep brain structures and their associated networks.

The unprecedented level of spatial precision achieved by our 256-element array, along with the integration of individualised treatment planning and closed-loop targeting, mark a significant advance over prior studies. Other recent work has demonstrated notable developments in transcranial ultrasound technology, enabling electronically-steered targeting of deep brain regions.^15,20^ However, these proof-of-concept studies focused on modulating the subgenual cingulate cortex, without demonstrating the ability to target specific thalamic nuclei. In contrast, our helmet transducer array achieves a focal volume nearly 30 times smaller than these systems, enabling selective targeting of structures as small as the lateral geniculate nucleus. This selectivity is reflected in the highly spatially-precise changes in BOLD activity in the visual cortex, as would be expected from precise targeting of the LGN.

Another recent study demonstrated the ability to induce a sustained reduction of essential tremor in patients by targeting the VIM and the DRT using a clinical MRgFUS system.^13^ While this work highlights the potential of highly-focal low-power, non-thermal ultrasound stimulation for therapeutic applications, the use of a neurosurgical frame and the need for focal tissue heating to confirm the target location limit its suitability for non-invasive neuromodulation studies in healthy participants. Our system overcomes these limitations by employing a custom-designed stereotactic face and neck mask for precise positioning and optimised model-based treatment planning and online re-planning for accurate targeting, enabling non-invasive studies in healthy populations. The advancements in spatial precision and targeting reliability open new avenues for studying the functional roles of specific deep brain nuclei and developing targeted, non-invasive therapies.

We chose to use a theta burst paradigm that has been widely used in the literature. The effects of this paradigm seem to vary across brain regions, with some studies demonstrating increased excitability and decreased inhibition when stimulating cortical regions,^16,17^ and others showing inhibitory after-effects.^21^ The mechanisms that result in this heterogeneity are not yet clear. It is plausible that differences in stimulation intensity in the stimulated region results in different effects; other non-invasive brain stimulation paradigms show substantial non-linearity in their response profiles,^22^ including a reversal from inhibitory to excitatory effects with increasing intensity.^23^ The significant effects of the stimulation on task-related activity were observed for at least 40 minutes after stimulation but were not still significant after 2 hours, consistent with previous findings using the theta burst protocol.^17,18^

Further research is needed to fully elucidate the mechanisms underpinning TUS and optimise stimulation parameters. A multiplicity of factors will likely influence the neuromodulatory effects of TUS, including the relative proportions of excitatory and inhibitory neurons within a target structure,^24^ the distribution of mechanosensitive cation channels along those neurons,^25,26^ and the morphology of the neuronal bodies and axons.^27^ The exact contribution of these factors has yet to be established for the range of ultrasound parameters used in vivo, but as our knowledge increases, these data can be used to optimise TUS parameters based on the unique properties of deep brain structures, potentially enhancing the efficacy and specificity of the technique.

To leverage the high spatial precision, we chose to use a dense-sampling approach, averaging across large amounts of data in a relatively small number of participants, rather than across spatially-varying stimulation targets across participants. This approach has proven highly effective in finely characterising neural circuits that are difficult to study with traditional scanning methods.^28–30^ However, the spatial focality of the TUS results in a very small volume of tissue within LGN being significantly stimulated. While this spatial precision represents a step-change advance in terms of specificity for neuromodulation, the size of the focus compared with the relatively large voxel size of our fMRI sequence means that we were unlikely to be sensitive to change in activity within the LGN itself, but rather observed the effects of stimulation in directly anatomically connected structures.

Several recent studies in animal models have demonstrated the ability of transcranial focused ultrasound stimulation to modulate activity in the visual thalamus (LGN) and produce downstream effects in the visual cortex. In sheep, reversible suppression of visually evoked potentials outlasting the stimulation period has been observed,^31^ while in mice, NMDAR-dependent long-term depression of thalamocortical synapses in the visual cortex has been reported.^32^ Furthermore, studies in non-human primates have shown sustained effects on visual choice behaviour and gamma activity following brief LGN stimulation.^33^ Our work builds upon these findings, demonstrating for the first time in humans that LGN-targeted ultrasound produces robust modulation of visual cortical activity, both during stimulation and enduring afterwards.

Our advanced TUS system enables selective and non-invasive modulation of deep brain structures in healthy adults. This approach holds promise for advancing human neuroscience beyond correlational understanding of deep structure function towards establishing causal relationships. This will allow the systematic probing of causal networks underlying the full range of cognitive processes on a participant-by-participant basis, something that had not previously been possible. TUS presents the tantalising possibility of extending our thorough understanding of the function of the deep structures in rodent models to humans. Furthermore, TUS offers the possibility of enhancing the specificity of existing therapeutic targets by elucidating the functional effects of stimulation prior to surgery, analogous to approaches that have already shown promise after surgical placement of DBS electrodes.^34^ The non-invasive nature of our approach facilitates this development, as TUS can be used to modulate multiple potential therapeutic targets in patients to assess likely behavioural effects before proceeding with invasive surgery. Furthermore, TUS itself could emerge as a standalone therapeutic option for certain disorders, offering a non-invasive alternative to surgical interventions.

In conclusion, our study presents a groundbreaking advance in deep brain neuromodulation, opening new avenues for both basic research and clinical applications. However, further work is needed to fully elucidate the mechanisms of action, optimise stimulation parameters, and establish the long-term safety and efficacy of this approach. Nonetheless, the unprecedented precision and non-invasive nature of our advanced TUS system hold immense potential for revolutionising our understanding and treatment of neurological and psychiatric disorders.

## Methods

### Transcranial ultrasound system

#### System overview

The MR-compatible transcranial ultrasound system is built around 256 individual transducer elements mounted within a semi-ellipsoidal helmet (Figure 1, Extended Data Figures 1-2). Each element is made from an air-backed piezocomposite material with an acoustic matching layer enclosed in a custom-designed plastic housing (Sonic Concepts). These elements, each with a 3 mm aperture diameter and operating at a centre frequency of 555 kHz, are connected via 40 cm micro-coaxial cables to 8 interconnect printed circuit boards (PCBs). The selected driving frequency provides a reasonable compromise between aberration and attenuation due to the skull bone and the size of the acoustic focus.^35,36^ The PCBs are connected to eight cables, each 8.2 metres long to allow them to reach from the inside of the MR bore to the MR penetration panel. Each cable contains 32 individually shielded twisted pairs within an overall copper braid designed to mitigate electrical cross-talk and capacitive coupling. The distal ends of these cables collectively terminate in two ultrasound connectors (DL5260P, ITT Cannon), and connect to a bespoke feed-through connector mounted on the MR feed-through panel, forming an RF bond. This connector also incorporates an electrical matching network to maximise power transfer. In the MR control room, a secondary set of cables links the feed-through connector to a Verasonics Vantage 256 ultrasound drive system.

#### Helmet dimensions

The shape of the semi-ellipsoidal helmet is designed to conform to the average adult head, based on an analysis of T1-weighted MR images from 16 healthy volunteers (ages 19-42 years, 11 female). Initially, we determined a suitable inclination angle for the helmet relative to a plane perpendicular to the scanner bed. We used the approximate positions of the air-filled frontal sinus and the external occipital protuberance as reference points, taking into account the participants’ head orientation in the scanner (Extended Data Figure 1a). This process yielded an optimal inclination angle of 20 degrees. We then determined the average head size by fitting an angled semi-ellipsoid to segmented head masks derived from the T1 images, resulting in average head dimensions of 206 mm in length, 157 mm in width, and 96 mm in height (Extended Data Figure 1b). To comfortably accommodate most adults while minimising the water volume and distance to the head, the helmet’s interior was designed to be 40 mm larger than these average dimensions.

#### Element layout

The transducer elements were randomly distributed across the helmet surface, with each element’s normal oriented towards the centre of the semi-ellipsoid. This random positioning strategy mitigates the formation of significant grating lobes, a potential concern due to the relatively low element count and the average spacing exceeding half the acoustic wavelength. To ensure line of sight to deep brain structures, we employed an offset angle of 15 degrees from the helmet’s lower exterior (Extended Data Figure 1c). Additionally, we limited the element arrangement to an upper segment angle of 55 degrees, chosen based on numerical simulations to balance focal size and side lobe height optimally (Extended Data Figure 1d). A minimum distance of 10 mm between elements was maintained for manufacturability, with additional exclusion zones to allow water connections at the highest and lowest points. The final element positions were determined from 5000 numerical simulations, selecting the configuration that minimised the relative sidelobe height (range 21 to 28%).

#### Helmet construction

The ultrasound helmet was fabricated using a HP-Jet Fusion printer, employing PA12 Nylon for its high mechanical strength (Extended Data Figure 1f). Each transducer element housing was specifically designed with a flange to facilitate accurate axial placement within designated apertures in the helmet (Extended Data Figure 2a). This modular approach not only ensured precise positioning but also allowed for the individual elements to be conveniently removed and replaced if necessary, enhancing the system’s maintenance and longevity. The elements were then securely coupled to the helmet using silicone adhesive. To accommodate the interconnect printed circuit boards (PCBs) and provide necessary strain relief for the cables, the helmet was integrated with a rigid enclosure (Extended Data Figure 2a). Additionally, a custom vacuum-formed liner was employed on the MRI bed, designed to prevent potential water spills from reaching the MRI bed, thereby safeguarding the integrity of the MRI system.

#### Driving system

The transcranial ultrasound system was operated using the Verasonics Vantage 256 Research Ultrasound System with an external power supply (HIFU configuration). System control and parameter adjustments were facilitated through a custom MATLAB graphical user interface (GUI). To mitigate the audibility of the ultrasound stimulation, pulse ramping was implemented using the ‘states’ transmit waveform type which allows repeated transmission of a tri-state waveform with a given amplitude.^37,38^ The ramp was implemented by discretizing a raised cosine ramp (Tukey window) into 50 linearly spaced amplitude levels, allowing for a gradual increase or decrease in ultrasound intensity over the ramp duration. For experiments involving concurrent functional magnetic resonance imaging (fMRI), we synchronised the ultrasound system’s triggering with the MR trigger out signal (Extended Data Figure 2b). This synchronisation enabled the interleaving of ultrasound stimulation with MR measurements, utilising a sparse acquisition approach to minimise interference between the two modalities.

#### Subject positioning system

For precise and repeatable positioning of participants within the helmet, we developed a custom-designed stereotactic face and neck mask, fabricated using 3D printing and casting techniques (Figure 1, Extended Data Figure 2a). The participant’s anatomical data, required for the mask’s design, was obtained from T1-weighted MR images captured using a 64-channel head coil. We extracted a skin mesh from these images to derive the mask’s structure. The mask comprised two parts: a neck support and a face mask, each designed to engage specific anatomical landmarks—the frontal, sphenoid, and temporal bones for the upper part, and the occipital bone and neck for the lower part—while leaving openings for the participants eyes, ears, and mouth. Quick-release connectors facilitated the secure assembly of these parts, and a flange on the neck support was used to locate the positioning system to the helmet. Both components were 3D printed using an Ultimaker S5 printer in PLA material. To improve comfort, the surface of the neck support and the face mask were covered with a cushioning layer, cast in soft silicone rubber (Ecoflex Gel 2), providing a conformal contact with the participant. Railings on each side of the face mask were incorporated to facilitate the attachment of an adjustable mirror, enabling participants to view the visual display unit positioned at the end of the MRI bore (Extended Data Figure 2).

#### Water coupling

Acoustic coupling between the ultrasound helmet and the participant’s head was achieved using deionized and degassed water, filling the space within the helmet. To facilitate effective coupling, participants’ heads were shaved to prevent air bubble entrapment. The water barrier was created using a flexible silicone membrane with a central hole, mounted within an ellipse-shaped sealing flange and clamped between the participant positioning system and the helmet (Figure 1, Extended Data Figure 2). The water was pre-heated to a physiological temperature of 37 degrees Celsius using a water conditioning unit (Sonic Concepts WCU-105) coupled to a 30 L external reservoir equipped with an additional internal heater. The helmet was filled through detachable hoses (10 metres long, 12 mm internal diameter) from its base, with an overflow chamber positioned at the top. The filling process was completed in approximately 3 minutes, while drainage took about 1 minute. After filling, the water was not circulated. Over the course of a typical 45-minute experiment, the water temperature inside the helmet was observed to decrease by approximately 4 degrees Celsius.

#### Air pressure control

The hydrostatic pressure on the participant’s head due to the water volume increases the contact force exerted by the positioning hardware, causing discomfort. To counteract this, we implemented a custom-built pneumatic controller to control the air pressure in the overflow chamber above the water, interconnected with the top of the helmet. This controller comprised a ported air pressure sensor with a digital readout and a diaphragm pump, both interfaced with a microprocessor. A control algorithm on the microprocessor maintained the air pressure in the chamber at a target value slightly below atmospheric pressure (set at 99500 Pa), effectively reducing the hydrostatic pressure on the participant’s head. The controller was situated in the MR control room, connected to the air chamber via a 12-metre-long tube with an internal diameter of 3 mm. This tube passed through a waveguide in the MRI penetration panel (Extended Data Figure 2).

#### System calibration

For accurate calibration of our ultrasound system’s acoustic output, we followed a systematic procedure. The helmet was affixed to an automated scanning tank equipped with a five-degree-of-freedom (x, y, z, rotate, tilt) scanning arm (Extended Data Figure 6a). This arm held a calibrated 200 µm needle hydrophone (Precision Acoustics) positioned at the array’s geometric centre, a location determined by the element positions and their time-of-flight measurements. We individually measured the signal from each element, extracting amplitude and phase at the driving frequency from the waveform’s steady-state segment. To standardise the acoustic output across elements, we calculated amplitude scaling factors and phase offsets for each, after accounting for the hydrophone’s directional response and the distance to each element. Amplitude scaling was refined through a calibration curve correlating the Verasonics apodization parameter (which controls the on-time of the tri-state pulser) with acoustic output amplitude. Finally, we established a relationship between driving voltage and pressure using the hydrophone’s calibrated sensitivity.

#### Acoustic performance

To assess the acoustic performance of our system, we conducted numerical simulations examining the acoustic output across a 16 cm steering range centred on the helmet’s geometric centre (Figure 2b). For each steering position, we evaluated the −3 dB focal size, focusing gain, and grating lobe height. A selection of these positions was experimentally validated using the same scanning setup as described in *System calibration*, employing a Fabry-Perot fibre optic hydrophone to minimise hydrophone directivity and spatial averaging effects. The excellent quantitative match between the simulated and experimentally measured amplitudes, profiles, and −3 dB beam sizes (Extended Data Figure 3) demonstrates the accuracy of the source definition used in numerical simulations and the near-ideal performance of the array. This close correspondence also indicates that any electrical or mechanical crosstalk is effectively negligible, thanks to the physical separation of the elements and the use of cables designed to eliminate the electrical crosstalk observed in a prototype system.^39^ At the geometric focus, the −3 dB focal size was measured at 1.3 mm laterally and 3.4 mm axially. This was relatively consistent across the lateral steering range (Figure 2b, Extended Data Figure 3b). The focal size decreased for positions inside the helmet and increased for those outside, reflecting changes in the effective aperture. Across a 5 cm steering range from the centre of the array (which covers the thalamus), the grating lobe height was less than 22%, demonstrating the effectiveness of the sparse random array design.

#### MR compatibility

To assess the impact of our ultrasound system on the MRI environment, we conducted an RF noise spectrum analysis using a Siemens 3T Prisma MRI scanner (Extended Data Figure 4a). This quality assurance scan measures the radiofrequency noise within the MRI scanner, which can be used to evaluate the system’s electromagnetic compatibility. The analysis was performed under several configurations to comprehensively evaluate potential noise sources. Adding our ultrasound helmet to the MRI, either water-filled or with a participant, elevated the thermal noise floor slightly due to the increased water volume in the receive coil’s field. Importantly, this did not result in any RF interference spikes, indicating the helmet’s compatibility with the MRI’s RF environment.

To evaluate the impact of our ultrasound helmet on MRI image quality, we conducted a second experiment comparing participant images acquired using the body coil, with and without the helmet (Extended Data Figure 4b). Analysis of these images revealed a contrast-to-noise ratio (CNR) of 30 in the reference case, compared to 26 when the helmet was used. This slight reduction in CNR indicates a modest impact of the helmet on image quality, a factor crucial for considering the helmet’s integration with MRI procedures. However, when the ultrasound system was activated during image acquisition, banded artefacts were observed in the MRI images. This is attributed to the incomplete RF shielding of the ultrasound system, leading to electromagnetic interference with the MRI’s signal. To mitigate this, during online Echo Planar Imaging (EPI) measurements, we employed a sparse imaging approach, interleaving MR measurements and ultrasound stimulation. This technique effectively prevented the artefacts by temporally separating the MRI acquisition from periods of ultrasound activity.

In a third experiment, we assessed the impact of the MRI environment on ultrasound transmission from the helmet (Extended Data Figure 4c). For this, we used the water-filled helmet with a blank cover, incorporating a Fabry-Perot fibre optic hydrophone positioned at its geometric centre. The helmet’s driving phases were adjusted to focus on the hydrophone. Acoustic waveforms were recorded from the hydrophone under various conditions: (1) with the system on the MRI bed but outside the bore, (2) inside the MRI bore with the scanner turned off, and (3) within the bore during the acquisition of a standard EPI MRI sequence. The analysis revealed negligible differences in the acquired waveforms across these conditions, with less than 1% variation in waveform amplitude. Additionally, when the hydrophone was used to passively monitor the environment while acquiring an EPI image, no discernible signals were detected beyond the inherent noise level. This indicates that the MRI environment does not affect the acoustic output of the helmet.

#### Subject positioning accuracy and repeatability

To assess the positioning repeatability of our custom-designed stereotactic face and neck mask across sessions, we performed pairwise comparisons of positioning images from three participants, acquired over multiple sessions. Each image, including the LGN target position, was registered to a defined helmet space (see *Treatment planning* for details). We then calculated the LGN positional differences across all images and sessions. The average shift in each Cartesian direction (in helmet coordinates) and the average overall shift (Euclidean distance) were computed. This gave average target shifts of 0.54 +/-0.43 mm and 0.48 +/-0.37 mm in the lateral directions, 1.13 +/-0.79 mm in the axial direction (into the helmet), and an overall average target shift of 1.50 +/-0.70 mm (mean+/-SD). These values demonstrate a high level of positioning repeatability, comparable to, and in some cases surpassing, the precision achieved by other stereotactic positioning systems used in similar neuroimaging contexts.^40,41^

To assess the efficacy of the face and neck mask in minimising head movement during MRI acquisition, participant motion was quantified through a motion correction process implemented in FEAT (FMRIB’s Software Library v6.0). Functional scans underwent realignment using rigid body transformations, enabling the calculation of six movement parameters (three translations and three rotations) for each scan, relative to a designated reference volume. During representative scans (all on-line, on-target scans for the first three participants), the mean participant movement was 0.25 +/-0.001mm (mean+/-SD), demonstrating a very high level of positioning stability within sessions.

### Treatment planning

#### Participant demographics

Seven healthy participants (6 male, age range 28-54) gave their written informed consent to participate, in line with ethical approval from the East Midlands Leicester South Research Ethics Committee (22/EM/0164). Participants had no history of neurological or psychiatric conditions and had normal or corrected-to-normal vision, and were not taking any psychoactive medications.

Exclusion criteria included contraindications to MRI and to non-invasive brain stimulation, including a personal or family history of seizures or epilepsy. The study was performed in accordance with the Declaration of Helsinki, except for pre-registration.

#### Study visits

The overall study was designed as two dense-sampling experiments on seven human participants split across seven visits (Extended Data Figure 5c). In visits 1 to 3, MR and CT planning images were obtained (see *Planning images*). In visits 4 and 5, we conducted the online TUS experiment. In visits 6 and 7, we conducted the offline TUS experiment. Stimulation sessions were spaced by at least 1 week. Of the seven participants, four completed all seven visits, two took part in visits 1-5 and one took part in visits 1-4.

#### Visual stimulus

To elicit functional activity in the LGN and visual cortex, we utilised a radial checkerboard pattern stimulus. Stimuli were displayed on a monitor positioned at the end of the MRI bore and viewed via mirrors mounted on the coil or helmet (Extended Data Figure 2). The stimulus, set at a 50% contrast level relative to total screen luminance and reversing contrast at 7.5 Hz, was designed to balance visual engagement with comfort. This pattern has been previously been shown to effectively stimulate both primary and secondary visual processing areas.^42,43^ While images acquired using the head and neck coil allowed for full visibility of the stimulus, the presence of the helmet restricted the view to a rainbow-shaped section passing through the centre of the checkerboard (Extended Data Figure 4d). However, this reduced visibility still proved sufficient to robustly elicit the desired neural activity in the targeted regions (Extended Data Figure 4e). During the task, the visual stimulus was displayed in 15-second blocks followed by a 9-second break during which a fixation cross was displayed. This was repeated 20 times, giving a total task time of 8 min 9 sec.

#### Planning images

For treatment planning, images were acquired across three sessions (Extended Data Figure 5a,c). In the first session, planning images were acquired using a Siemens 3-T Prisma whole-body MRI scanner with a 64-channel head and neck receive coil as follows:

- Large field-of-view (head and neck) T1-weighted MPRAGE for creation of the head and neck masks [voxel size 1 × 1 × 1 mm, repetition time (TR)=2300 ms, inversion time (TI)=900 ms, echo time (TE)=2.28 ms, field of view (FOV)=192 × 288 × 288 mm, sagittal, flip angle(FA)=8°, PAT factor = 2, no fat saturation, total acquisition time (TA)=5:58 (minutes:seconds)].
- High SNR T1-weighted MPRAGE scan optimised for grey and white matter contrast (voxel size 1 × 1 × 1 mm, axial, TR=1900 ms, TE=3.96 ms, flip angle=8°, TI=912ms, FOV = 232 x 256 x 192mm, no PAT, fat suppressed, TA=7:21).
- Resting state fMRI scan acquired with a T2*-weighted 2D multiband EPI sequence [voxel size 2.4 × 2.4 × 2.4 mm, TR=735 ms, TE=39.00 ms, FOV=210 × 210 × 154 mm, FA=52°, TA =6:10, multi-band acceleration factor (MB)=8].^62^ Participants were asked to fixate on a white cross presented on a black screen, to blink normally, and to try not to fall asleep.
- Diffusion [voxel size 2 × 2 × 2 mm, TR=3600 ms, TE=92.00 ms, FOV=210 × 210 × 144 mm, FA=78°, TA=6:32, MB=3, 100 diffusion encoding directions on two shells (b = 1000 s/mm2 and b=2000 s/mm2), and four b=0 images]. This was followed by a set of three b=0 images with reversed phase encoding direction to allow for distortion correction. Participants watched a nature video during this scan.
- Task fMRI scan acquired with a T2*-weighted 2D multiband EPI sequence (see *Visual stimulus*) (voxel size 2.0 × 2.0 × 2.0 mm, TR=1500 ms, TE=25 ms, FOV=216 × 216 × 144 mm, FA=70°, TA=8:30, MB=3).
- B0 field map to enable distortion correction in the BOLD fMRI data (voxel size 2.5 × 2.5 × 2.5 mm, TR=590 ms, TE1=4.92 ms, TE2=7.38 ms, FOV=210 × 210 ×150 mm, FA=46°, TA=1:40).

The T1-weighted structural images and the single-band reference images from the T2*-weighted BOLD scans were pre-processed, bias-corrected and brain-extracted using `fsl_anat` and `BET` tools from the FMRIB Software Library (FSL).^44,45^ Fieldmaps were processed and brain extracted using `BET` and `fsl_prepare_fieldmap` FSL tools. The task fMRI data was processed and registered to MNI standard space through first-level `FEAT` analysis.^46^ The 4D data from each task run was modelled using a general linear model (GLM) in lower-level `FEAT` with one explanatory variable (EV) modelling the checkerboard presentation blocks.

In the second session, a low-dose CT scan was acquired using a GE Revolution CT scanner to obtain the participant’s bony anatomy, essential for treatment planning simulations [pixel spacing=0.45mm (typical value), slice thickness: 0.625 mm, convolution kernel: BONEPLUS, tube current: 70 (typical value), KVP: 80]. An electron density phantom (CIRS Model 062M) was also scanned using the same acquisition and reconstruction parameters to allow a precise calibration from Hounsfield units (HU) to mass density to be determined.^47^

In the third session (and during the online stimulation sessions), participants were positioned within the water-filled helmet and images were acquired using a Siemens 3-T Prisma whole-body MRI scanner with the body coil. Before the scans were acquired, the shim volume was set manually to cover the brain but exclude the water in the helmet, and the GRE Brain shimming routine was manually iterated three times, followed by manual frequency adjustment. The initial manual shim was then applied to the following scans:

- T1-weighted magnetisation prepared (MPRAGE) structural scan (voxel size 1.5 × 1.5 × 1.5 mm, TR=1690 ms, TE=3.78 ms, TI=904 ms, FOV=306 × 336 × 264 mm, FA=8°, TA=5:45).
- A repeat of the MPRAGE with the inversion pulse disabled to maximise signal from the water in the ultrasound helmet for registration.
- Task fMRI scan (see *Visual stimulus*) acquired with a sparse T2*-weighted 2D EPI sequence (voxel size 3 × 3 × 3 mm, TR=3000 ms, volume TA=2600 ms, resulting in a 400 ms idle period in each repetition, TE=29.0 ms, FOV=300 × 300 × 114 mm, FA=90°, TA=8:14).
- B0 field map to enable distortion correction in the BOLD fMRI data (voxel size 4.2 × 4.2 × 6 mm, TR=440 ms, TE1=4.92 ms, TE2=7.38 ms, FA=45°, TA=1:11).

#### Target identification

To determine the target voxel for the lateral geniculate nucleus (LGN), we integrated data from three sources. First, functional MRI (fMRI) data acquired during the visual task in the planning session were processed as described above and then the site of maximum activation within the thalamus was identified. Second, we utilised a high-resolution LGN atlas,^48^ which was transformed into participant space by non-linearly registering the participant’s planning image with the MNI head template. Third, the thalamic nuclei were segmented using FreeSurfer, generating a parcellation of the thalamus into distinct nuclei based on a probabilistic atlas derived from histological data.^49^ Based on this composite information, we manually selected the 1×1×1 mm LGN voxel that exhibited the highest activation within the bilateral functional mask and overlapped with both structural LGN masks (Figure 2c). For the active control site, our target voxel was chosen in the magnocellular medial dorsal nucleus (MDN) as identified by the FreeSurfer segmentation. Targets were selected in the right hemisphere for participants 1, 4, 5 and 7, and the left hemisphere for participants 2, 3, and 6. For each participant, we selected the LGN with the most robust visually-evoked activity in the planning session. The control location was then selected in the same hemisphere as the active target. All data from participants with targets in the right hemisphere was flipped prior to group analysis, so that the stimulated LGN always appears in the left hemisphere for group analyses.

#### Offline planning

To map between helmet, brain, and CT coordinates, the acquired planning and positioning images underwent a three-step registration process (Extended Data Figure 5b). Initially, the positioning image was brain-extracted using FSL’s `BET` tool.^44^ The positioning image with the brain removed was then registered to helmet space, aligning it with a water reference image derived from CAD drawings of the helmet. Subsequently, the extracted brain images from both positioning and planning sessions were registered. Finally, the CT image, resampled to isotropic resolution using trilinear interpolation, was registered to the high-resolution planning image. This allowed the helmet position and the target positions to be mapped to the CT image coordinates for running planning simulations (Figure 2c). All registrations were performed using FSL’s `FLIRT` with six degrees of freedom and a mutual information cost function.^50–52^

Model-based treatment planning for our ultrasound system was conducted using k-Plan, our commercially available treatment planning software. This software computes the requisite driving amplitude and phase for each transducer element to focus the acoustic array on the specified target position with the desired target pressure. It also predicts the resulting acoustic pressure and temperature fields within the head. These calculations are based on a full-wave acoustic model which accounts for the unique acoustic properties of the skull and brain tissues. A custom transducer model, matching our physical transducer’s specifications, was integrated into k-Plan. Acoustic properties for each participant were mapped from low-dose CT scans, incorporating the CT calibration data. Simulations were performed with a resolution of 8 points per wavelength (Extended Data Figure 5d).^53^

The target was selected as described in *Target identification* and the target acoustic pressure for all experiments and participants was set to 775 kPa. This corresponds to 20 W/cm^2^ pulse average intensity assuming a characteristic impedance of 1.5 MRayls. In [X, Y, Z] helmet coordinates (see Extended Data Figure 2a), the mean position of the left LGN was at [-9, 23, −13] mm, the mean position of the MDN was [2, 1, −5] mm, and the mean distance from the LGN to the MDN was 23 mm. Across all participants and targets, the simulations predicted an in situ pressure amplitude at the target of 775 kPa, a maximum temperature increase in the brain of less than 0.2 degrees Celsius, and a mechanical index MI_TC_ of less than 1.9, in accordance with the ITRUSST consensus on biophysical safety.^54^ The focal size in water at the average LGN target position in [X, Y, Z] helmet coordinates was: [1.4, 1.6, 1.8] mm (−3 dB free-field), [2.0, 2.3, 2.6] mm (−6 dB free-field), while the average focal size in situ was [1.5, 1.5, 2.2] mm (−3 dB in situ), and [2.2, 2.2, 3.1] mm (−6 dB in situ) (parameter summary included in Extended Data Figure 7).^55^ Note the helmet axis are not necessarily aligned with focal ellipsoid.

#### On-line re-planning

To accommodate small changes in participant position relative to the helmet between sessions, we implemented a re-planning protocol, crucial for maintaining precision in targeting. Given that the ultrasound focal size, thalamic target size, and potential participant shifts are all in the 1-5 mm range, precise alignment is essential. At the beginning of each stimulation session, an additional positioning image was acquired to determine any target shift in helmet coordinates, following the same methodology as outlined in *Subject positioning accuracy and repeatability* (Extended Data Figure 5b). The driving phases for each helmet element, initially calculated using k-Plan, were then adjusted by adding geometrically calculated phase offsets (Extended Data Figure 5b). These adjustments aimed to shift the acoustic focus to align with the desired target position, leveraging the concept of the isoplanatic angle. This concept, borrowed from astronomy, posits that within a certain range, small shifts in participant position do not significantly alter phase distortions through the skull, thus allowing geometric adjustments to the initial phase calculations for accurate targeting.^56,57^ The advantage of the geometric refocusing is that it can be computed in real time, while the k-Plan simulation must be computed offline. Across all participants and all online stimulation sessions, the maximum shift required was 3.0 mm, with an average value of 1.86 +/-0.56 mm.

#### Experimental validation

To validate our treatment planning workflow, we conducted experiments using five human skulls, previously sectioned along a transverse plane above the ear line to isolate the cranial portion (Extended Data Figure 6). The skulls were obtained under a material transfer agreement in accordance with the UK Human Tissue Act. Prior to each experiment, these skull caps were submerged in deionized water, degassed at −400 mbar for 48 hours, and air-dried post-use. After CT scanning the skull caps, we extracted a surface mesh from the scans to design and 3D print mounts, securing the skulls in a known position relative to the helmet and the scanning tank. For each skull, we executed treatment plans targeting four positions: the geometric centre of the array and three points offset by 20 mm in each Cartesian direction within helmet coordinates. The experiments involved driving the array with a 45-cycle quasi-continuous wave signal, with phases determined by the treatment plan. Acoustic measurements were conducted in the previously described measurement tank using a fibre-optic hydrophone (FOPH), where we acquired line scans through the location of spatial peak pressure (Extended Data FIgure 6d). These measurements were processed to determine the focal size, amplitude, and position, and then compared to the planned values. The results showed that on average the measured spatial peak pressure values were within 21% of the target pressure, and the focal position within 0.9 mm. The mean −3dB focal dimensions were (x, y, z) = (1.3, 1.5, 3.1) mm (Extended Data Figure 6f), with an average difference of (dx, dy, dz) = (0.2, 0.2, 0.7) mm from the planned −3dB focal dimensions, and (dx, dy, dz) = (0.1, 0.2, 0.6) mm from the corresponding −3dB free field focal dimensions. These findings confirm the accuracy and precision of our helmet array and the associated treatment planning software and workflow.

In an additional experiment on one skull, we validated our re-planning protocol by geometrically shifting the focus by 5 mm from two planned positions, and again acquiring line scans through the location of spatial peak pressure. The results showed that on average the measured spatial peak pressure values were within 12.5% of the unshifted values, the focal positions within 0.2 mm of the intended shift position, and the differences in −3dB focal dimensions from the planned positions were (dx, dy, dz) = 0.03, 0.2, 0.8 mm (Extended Data Figure 6g). These findings validate the precision of our re-planning method and confirm the isoplanatic assumption for small positional adjustments.

### Online Stimulation of LGN and Visual Cortex Activity

#### Experimental design

We employed a single-blind, pseudo-randomized, sham-controlled block design to investigate the effects of transcranial ultrasound stimulation (TUS) on the visual system. During the experiment, participants fixated on a central point while a visual checkerboard stimulus was presented (see *Visual stimulus*). Each session consisted of 20 blocks, each lasting 15 seconds, during which the visual stimulus was displayed. TUS was applied during 10 of these blocks, while the other 10 blocks served as sham stimulation, with the order of active and sham blocks pseudo-randomized and balanced within each session. The block design was kept consistent within and across participants. Active TUS was delivered in 300 ms pulses every 3 seconds during active blocks (TUS was synchronised with the block timing, but not the individual checkerboard reversals), with a 10 ms ramp-up and ramp-down period to minimise auditory artefacts (see *Driving system*). The operator of the TUS system was unblinded due to audible and visual indicators from the control system when the TUS is active. Functional MRI (fMRI) measurements were acquired every 3 seconds, interleaved with the TUS pulses (see *Planning images*). Each participant underwent two online stimulation days, each including up to six MRI sessions, except for one participant who only took part in one online stimulation day. The number of stimulation runs per session varied between one and four, depending on the participant’s comfort level, with a maximum of six on-target stimulation runs per day. For three participants, three additional off-target stimulation runs were conducted, with the TUS focus targeted at an active control site adjacent to the LGN (described in *Target identification*). This experimental design allowed for a robust comparison of the effects of active TUS versus sham stimulation on both the targeted deep brain structure (LGN) and its functionally connected cortical region (V1) while controlling for potential confounds.

#### Magnetic resonance imaging

An identical MRI approach was used for the TUS sessions as during the third planning session (see *Planning images*), except that more than one task sequence was performed when the participant was comfortable enough to continue with the scan.

#### Image processing

T1-weighted images were brain-extracted and bias-corrected using the `BET` and `FAST` tools from FSL.^44,58^ Manual adjustments were applied to the brain extraction process when `BET` could not entirely eliminate water. Fieldmaps were similarly brain-extracted using `BET`, followed by preprocessing with the `fsl_prepare_fieldmap` tool. To address minor spiking artefacts which occurred on some EPI measurements when using body coil imaging (these were unrelated to the presence of the helmet, and eventually resolved by the scanner manufacturer), images were preprocessed using `MELODIC` independent component analysis (ICA), including motion correction using `MCFLIRT`, B0 fieldmap unwarping, high-pass temporal filtering at 100 seconds, and no spatial smoothing.^59^ Spiking artefacts were manually identified based on their spectral profiles—characterised by high power in a single volume—and removed using the `fsl_regfilt` tool. The resulting data were further processed through `MELODIC` ICA after applying a 5 mm FWHM smoothing filter. Components were then automatically labelled and noise components were cleaned using `FIX`.^60,61^ The cleaned data were processed and registered to MNI standard space via first-level `FEAT` analysis.^46^ For four participants, data were flipped after cleaning but prior to statistical analysis using the `fsl_swap_dim` tool to ensure that the stimulated LGN appeared in the left hemisphere across all seven participants.

#### Statistical analysis

The 4D data from each task run was modelled using a general linear model (GLM) in first-level `FEAT` with two explanatory variables (EVs) representing the active and sham blocks, respectively, and contrasts comparing brain activity between these blocks. Higher-level FEAT analysis was conducted using mixed effects (FLAME 1+2) for group analysis between participants, with separate EVs for each participant. The task mean activity map was employed as a pre-threshold mask, and a mixed effects analysis was employed with an automatic outlier de-weighting and a cluster correction of Z=3.1 and a p threshold of 0.05. A similar analysis plan was used for the three off-target runs of three participants.

### Offline Stimulation of LGN and Visual Cortex Activity

#### Experimental design

We employed an unblinded design with an active control site to investigate the long-lasting effects of transcranial ultrasound stimulation (TUS) on the visual system. Four participants underwent two offline stimulation sessions, one targeting the lateral geniculate nucleus (LGN) and the other targeting an active control site (MDN). To measure brain response, participants underwent three functional magnetic resonance imaging (fMRI) scans: a baseline measurement before stimulation (maximum 1 hour between the end of the scan and the stimulation), two early post-stimulation task scans (scanning started 19-21 minutes after stimulation and each scan lasted 8 minutes), and two late post-stimulation scans (scanning started approximately 140 minutes after stimulation and each scan lasted 8 minutes). Therefore, task fMRI data was collected on average between 20-40 minutes after stimulation for the early scan and between 140-160 minutes after stimulation for the late scan. During the stimulation session, participants fixated on a blank white screen while TUS was applied using a theta burst protocol. The stimulation lasted for a total of 80 seconds, with 20 ms of stimulation repeated every 200 ms (pulse repetition frequency: 5 Hz; duty cycle: 10%; 1 ms ramp-up and ramp-down). This design allowed for the assessment of both immediate and prolonged effects of TUS on visually-evoked brain activity, while the active control site served to demonstrate the specificity of the stimulation effects.

#### Magnetic resonance imaging

Participants had three MRI scans (baseline, early, late) acquired using a Siemens 3-T Prisma whole-body MRI scanner with a 64-channel head and neck receive coil as outlined below. The three scans were identical, except the T1 was acquired first at baseline and last in the early and late scans.

- T1-weighted MPRAGE scan acquired in the axial plane (voxel size 1 × 1 × 1 mm, TR=1900 ms, TE=3.96 ms, FA=8, FOV=232 × 256 × 192 mm, TI=912ms, TA=7:21).
- Resting state fMRI scan acquired with a T2*-weighted 2D multiband EPI sequence (voxel size 2.4 × 2.4 × 2.4 mm, TR=735 ms, TE=39.00 ms, FOV=10 × 210 × 154 mm, FA=52°, TA=10:00, MB=8).^62^ Participants were asked to fixate on a white cross presented on a black screen, to blink normally, and to try not to fall asleep.
- Task fMRI scan acquired with a T2*-weighted 2D multiband EPI sequence (see *Visual stimulus*) (voxel size 2.4 × 2.4 × 2.4 mm, TR=1500 ms, TE=32.40 ms, FOV=210 × 210 × 154 mm, FA=70°, TA =8:12, MB=4).^62^ Most MRI scans had 2 task runs, except for participant 2 late scan, which only had 1 task run (both on- and off-target).
- B0 field map to correct for distortion in the BOLD fMRI data (voxel size 2.5 × 2.5 × 2.5 mm, TR=590 ms, TE1=4.92 ms, TE2=7.38 ms, FOV=210 × 210 × 150 mm, FA=46°, TA=1:40).

#### Image processing

T1-weighted structural images and the single-band reference images from each T2*-weighted BOLD scan were pre-processed, bias-corrected, and brain-extracted using the `fsl_anat` and `BET` tools from FSL. Fieldmaps were processed and brain-extracted using the bet and `fieldmap_prepare` FSL tools. Exploratory analysis and preprocessing of the 4D task scans were performed using `MELODIC` ICA, which included motion correction using `MCFLIRT`, B0 field map unwarping, high-pass temporal filtering at 100 seconds, and no spatial smoothing. The `FIX` tool was then used to automatically label and clean the ICA components. The cleaned data was processed and registered to the MNI standard space through first-level `FEAT` analysis. For one participant, the data was flipped after cleaning, but before running the statistical analysis, using the `fsl_swap_dim` tool. This step was performed to ensure that the stimulated LGN appeared in the left hemisphere for all four participants, facilitating group-level analyses and comparisons.

#### Statistical analysis

The 4D data from each task run was modelled using a GLM implemented in `FEAT`. The GLM included one EV modelling the checkerboard presentation blocks. Higher-level FEAT analyses were conducted using a mixed-effects model (FLAME 1+2) for repeated-measures between-participant group analysis across time, performed separately for each stimulation target (LGN and active control site). For this group analysis, we used the task mean cortical activation map as a pre-threshold mask and automatic outlier-deweighting. Significant clusters were identified using a z-threshold of 3.1 and a corrected cluster significance threshold of p=0.05. This approach allowed for the identification of brain regions showing significant changes in visually-evoked activity following TUS, while accounting for both within-participant and between-participant variability.

## Data Availability

Individual fMRI statistical outputs mapped to standard space, and acoustic measurement and simulation data will be made publicly available before publication of the manuscript.

## Code Availability

Image and data processing pipelines will be made publicly available before publication of the manuscript.

## Acknowledgements

We thank James Robertson, Adam Thomas, Silvia Schievano, Sierra Bonilla, Edward Zhang, David Marsh, Antonio Stanziola, Felix Lucka, Jiri Jaros, Marta Jaros, Nick Everdell, Kyle Morrison, and Polytimi Frangou for technical assistance. We thank the WIN radiographers, the Oxford Radiology Research Unit Team at Churchill Hospital, the Radiology Department at Great Ormond Street Hospital, Mohamed Tachrount, Aaron Hess, Howell Fu, and Will Clarke for imaging assistance. We thank Saad Jbabdi for analysis assistance.

This study was supported by the Engineering and Physical Sciences Research Council, UK (EP/P008860/1, EP/P008712/1, EP/S026371/1). The k-Plan simulations were supported by the Ministry of Education, Youth and Sports of the Czech Republic through the e-INFRA CZ (ID:90254). EM was supported by a UKRI Future Leaders Fellowship (MR/T019166/1) and in part by the Wellcome/EPSRC Centre for Interventional and Surgical Sciences (WEISS) (203145Z/16/Z). CJS was supported by a Wellcome Trust Senior Research Fellowship (224430/Z/21/Z). The Wellcome Centre for Integrative Neuroimaging is supported by core funding from the Wellcome Trust (203139/Z/16/Z and 203139/A/16/Z). This study was supported by the NIHR Oxford Health Biomedical Research Centre (NIHR203316). The views expressed are those of the authors and not necessarily those of the NHS, NIHR or the Department of Health. For the purpose of open access, the author has applied a CC BY public copyright licence to any Author Accepted Manuscript version arising from this submission.

## Author information

These authors contributed equally: Eleanor Martin, Morgan Roberts, Ioana F Grigoras, Olivia Wright, Tulika Nandi.

These authors jointly supervised this work: Bradley Treeby and Charlotte Stagg.

## Contributions

EM, MR, OW, and BET designed, built, and characterised the ultrasound system. TN, CJS, and IFG designed the study. TN, CJS, and BET obtained ethics approval. IFG recruited the participants. IFG, CJS, EM, MR, BET, TDB, and BTC acquired the data. IFG, CJS, and TDB analysed the neuroimaging data. SWR and JC optimised the imaging protocols and conducted MRI compatibility and RF shielding tests. BET, CJS, EM, and BTC secured funding. BET, CJS, IFG, EM, and MR wrote the manuscript. All authors contributed to editing.

---

**Extended Data Figure 1:**
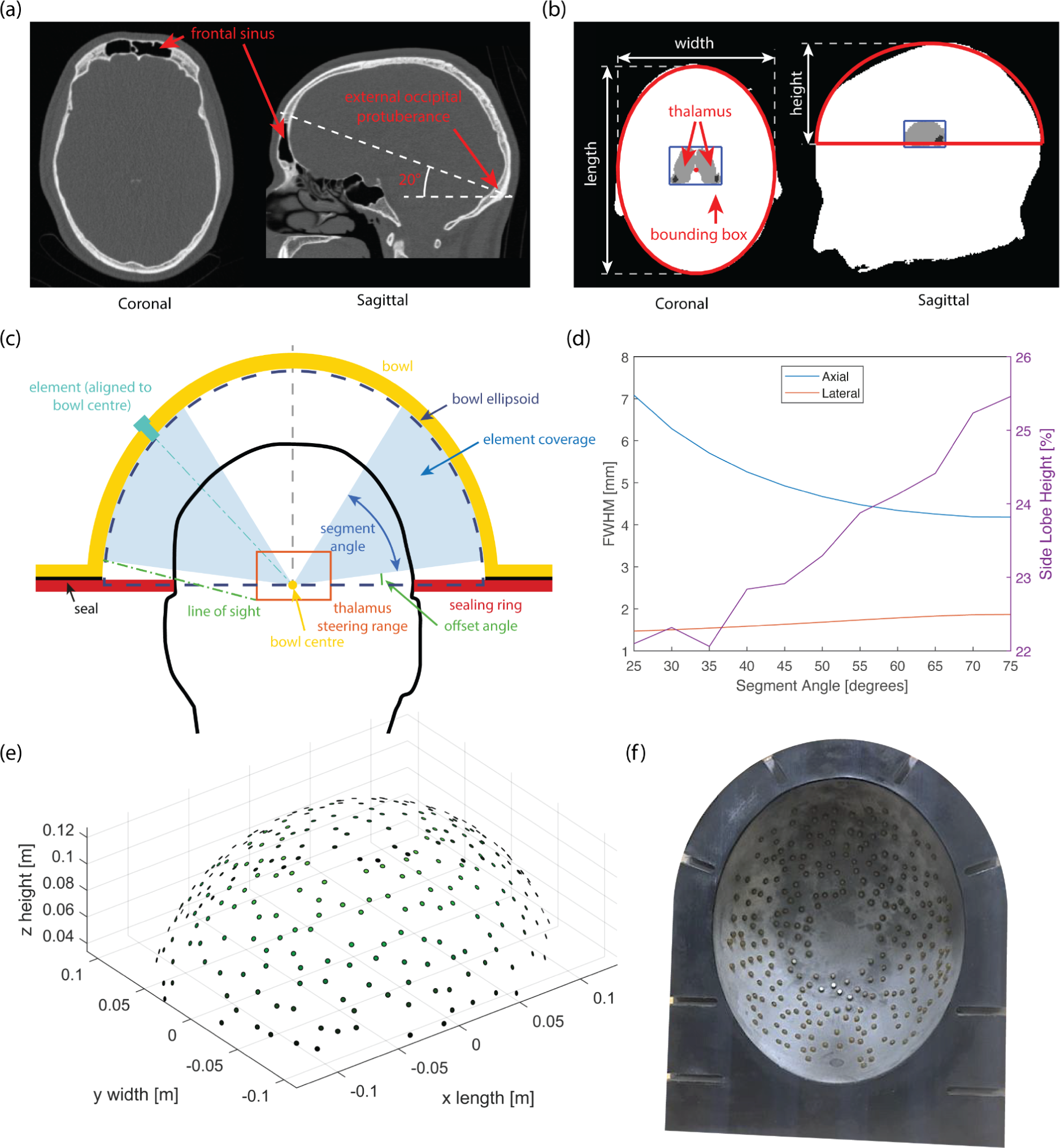
The helmet size was based on analysis of images from 16 healthy participants taking into account their head orientation in the scanner. **(a)** Illustrative CT image showing the chosen inclination angle of 20° and the approximate positions of the frontal sinus and external occipital protuberance for one participant. **(b)** Head size was determined for each participant by fitting an ellipse. The average ellipse dimensions were 206 × 157 × 96 mm (length × width × height). **(c)** The elements were selected to only cover a subset of the helmet ensuring line of sight to the average thalamus bounding box. **(d)** The segment angle was chosen to balance focal size with side lobe height. **(e)-(f)** Final element positions. The elements are randomly distributed on an ellipsoidal bowl with dimensions 286 × 237 × 135 mm (40 mm offset from average head ellipsoid), with a segment angle of 55°, and an offset angle of 15°. Two exclusion zones are also used to allow water connections at the highest and lowest points on the bowl.

**Extended Data Figure 2:**
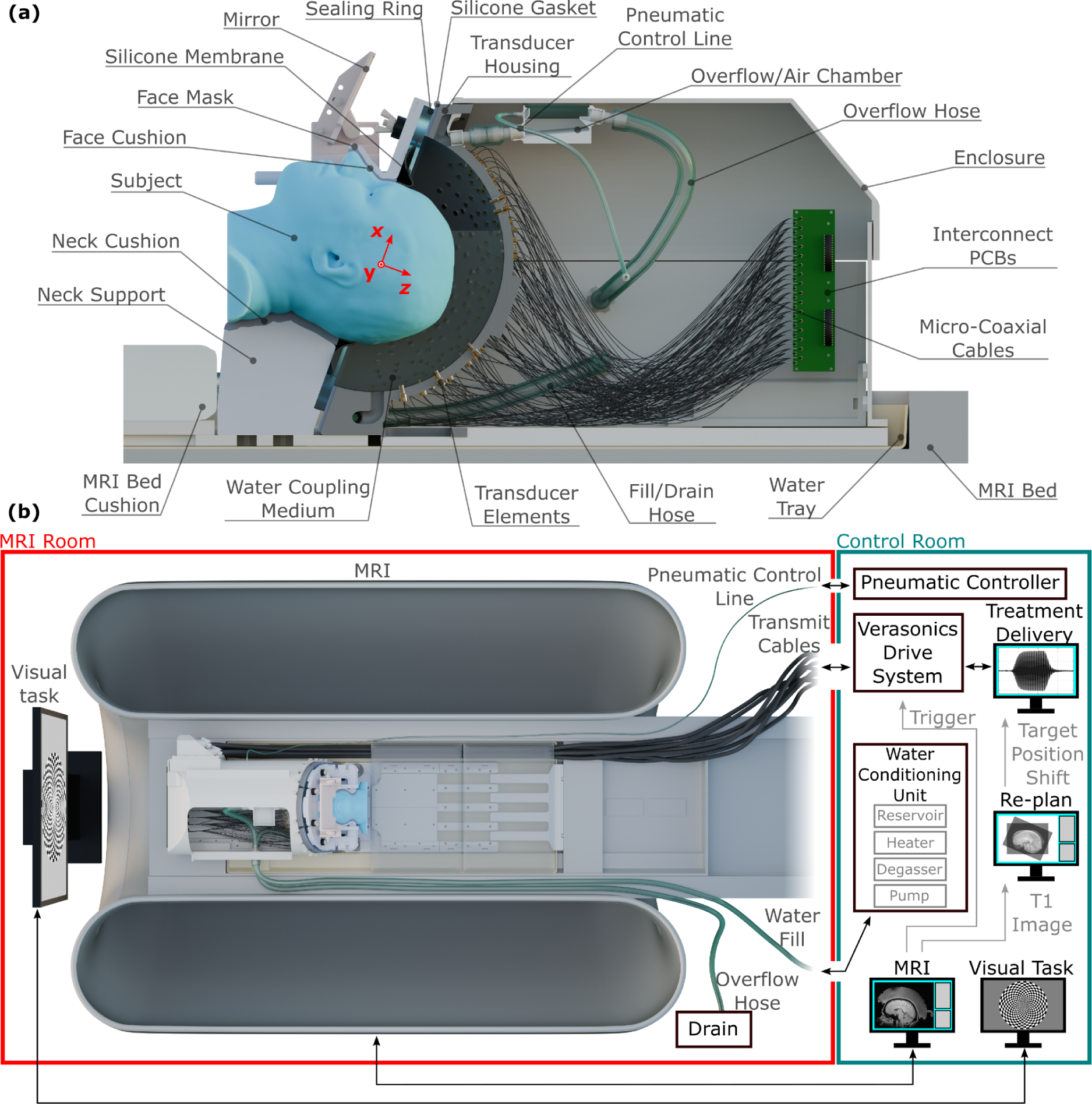
**(a)** Cross sectional view of the advanced transcranial ultrasound system. The participant is coupled to the transducer elements via a water layer, which is retained using a silicone membrane around the participant’s head. A gasket around the transducer housing flange prevents leakage. The transmit signals are routed via interconnect PCBs to the individual transducer elements. **(b)** Schematic view of the transcranial ultrasound system installed in the MRI, allowing concurrent neuromodulation and functional neuroimaging. Supporting equipment is located in the MRI control room. The water conditioning unit supplies water to the system via hoses. The transducer elements are connected to the Verasonics via 8 m cables. A pneumatic controller is used to restrict the hydrostatic pressure on the participant’s head, via an airline.

**Extended Data Figure 3:**
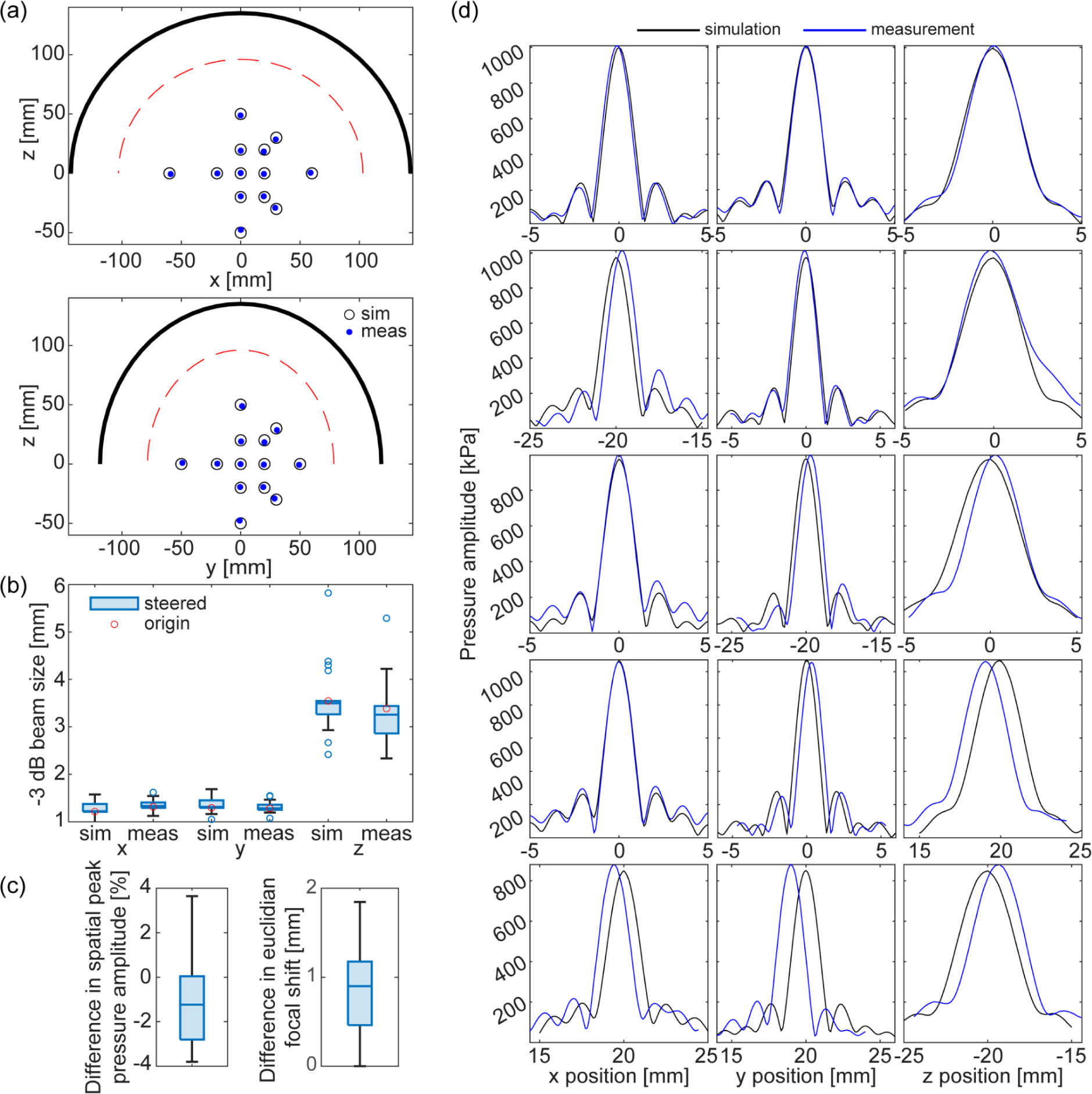
**(a)** Locations of steered free field validation measurements in the xz (top) and yz (bottom) planes. The bowl surface is shown in black and the average head by the red dashed line. Simulated focal positions using phases calculated geometrically are shown by black circles, blue dots show the position of the corresponding measured field steered using the same geometric phases. Measurement locations span the brain for the average head size. **(b)** Median and range (25th - 75th percentile) of simulated and measured free field −3 dB beam dimensions across 13 steering positions. Red circles show the focal dimensions at the origin, blue circles show outliers. Focal dimensions are preserved across the steering range. **(c)** Median difference between measured and simulated spatial peak pressure amplitude was −1.23%, and the difference was less than 4% across all positions. This demonstrates the accuracy of the transducer definition used in simulation, and the close to ideal performance of the array. Difference in focal shift is calculated from the absolute difference between the intended shift and the euclidean distance from the origin to the measured focal position. The median difference was 0.9 mm, and the difference was less than 2 mm across all positions. **(d)** Simulated (black) and measured (blue) pressure amplitude profiles through the focus in water at the origin, (0, 0, 0) mm, and for steering locations (−20, 0, 0) mm, (0, −20, 0) mm, (0, 0, 20) mm and (20, 20, −20) mm. A drive voltage of 8 V was used for all measurements, resulting in a spatial peak pressure of 1 MPa at the origin.

**Extended Data Figure 4:**
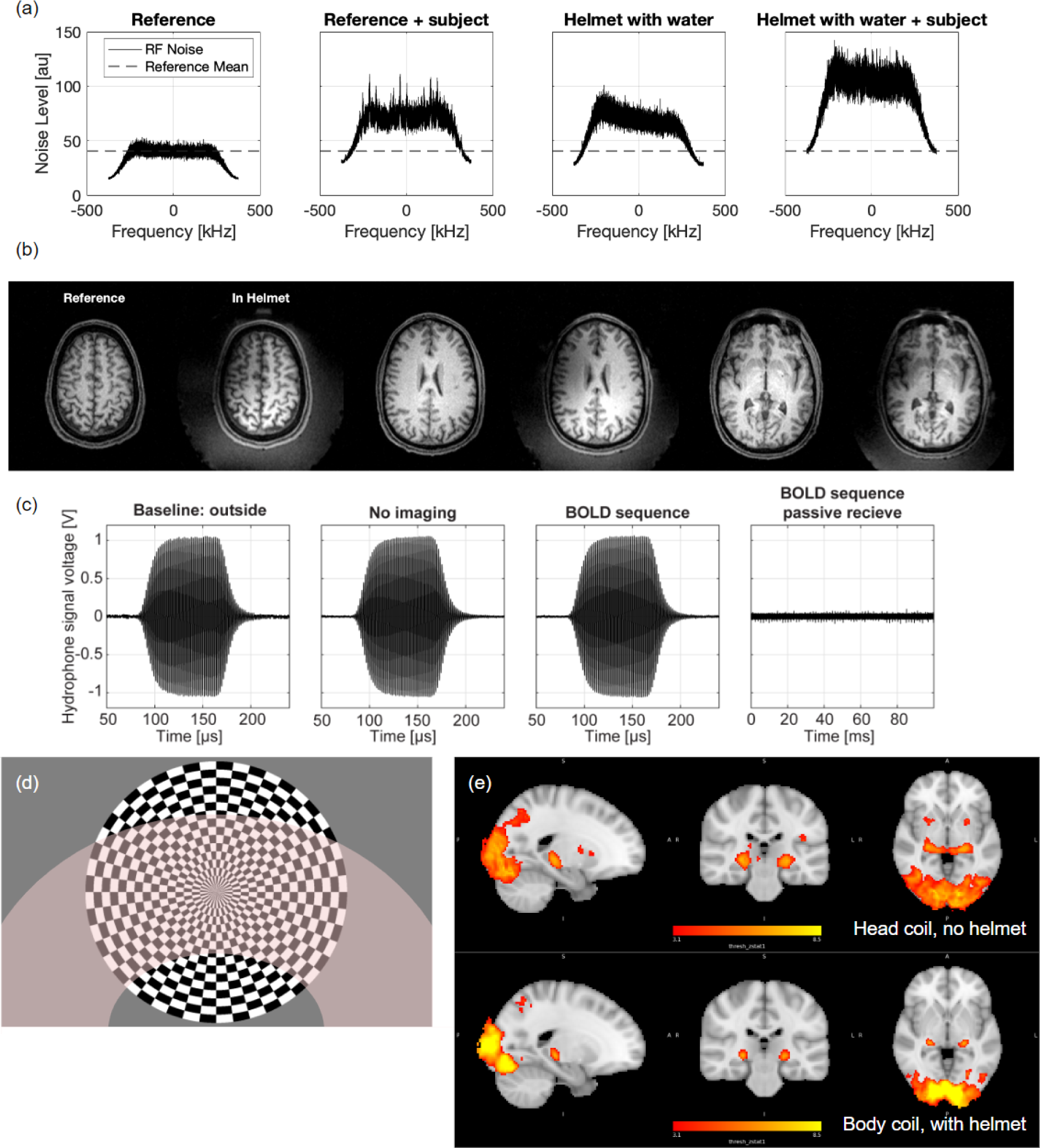
**(a)** Radio-frequency (RF) noise spectrum analysis of the MR system in various states with and without the helmet, and with and without the participant demonstrating good compatibility with the MR environment. **(b)** Pairs of body coil images (coronal slices) of the same participant with and without the water-filled helmet present. **(c)** Comparison of fibre-optic hydrophone measurements of the focal waveform acquired with 2 averages at 8V drive voltage, outside the MR bore, inside the MR bore in the absence of any imaging sequence, and inside the bore during the BOLD imaging sequence shows no change in acoustic waveform amplitude. Recording hydrophone voltage over a longer period (1 average) detected noise spikes which were small compared to the transmitted waveform amplitude. The noise equivalent voltage calculated as 3 standard deviations of the voltage in the absence of the acoustic signal was 15.5 ± 4 mV on average and did not significantly differ for the in bore recordings. **(d)** Radial checkerboard visual stimulus used to elicit functional activity in the LGN and visual cortex. The approximate field of view when the helmet is present is shown with the shaded area. **(e)** Mean activation during visual task using a standard 64-channel head and neck coil (top) and using the body coil with the helmet present (bottom) showing robust measurements despite the reduced field of view and sensitivity.

**Extended Data Figure 5:**
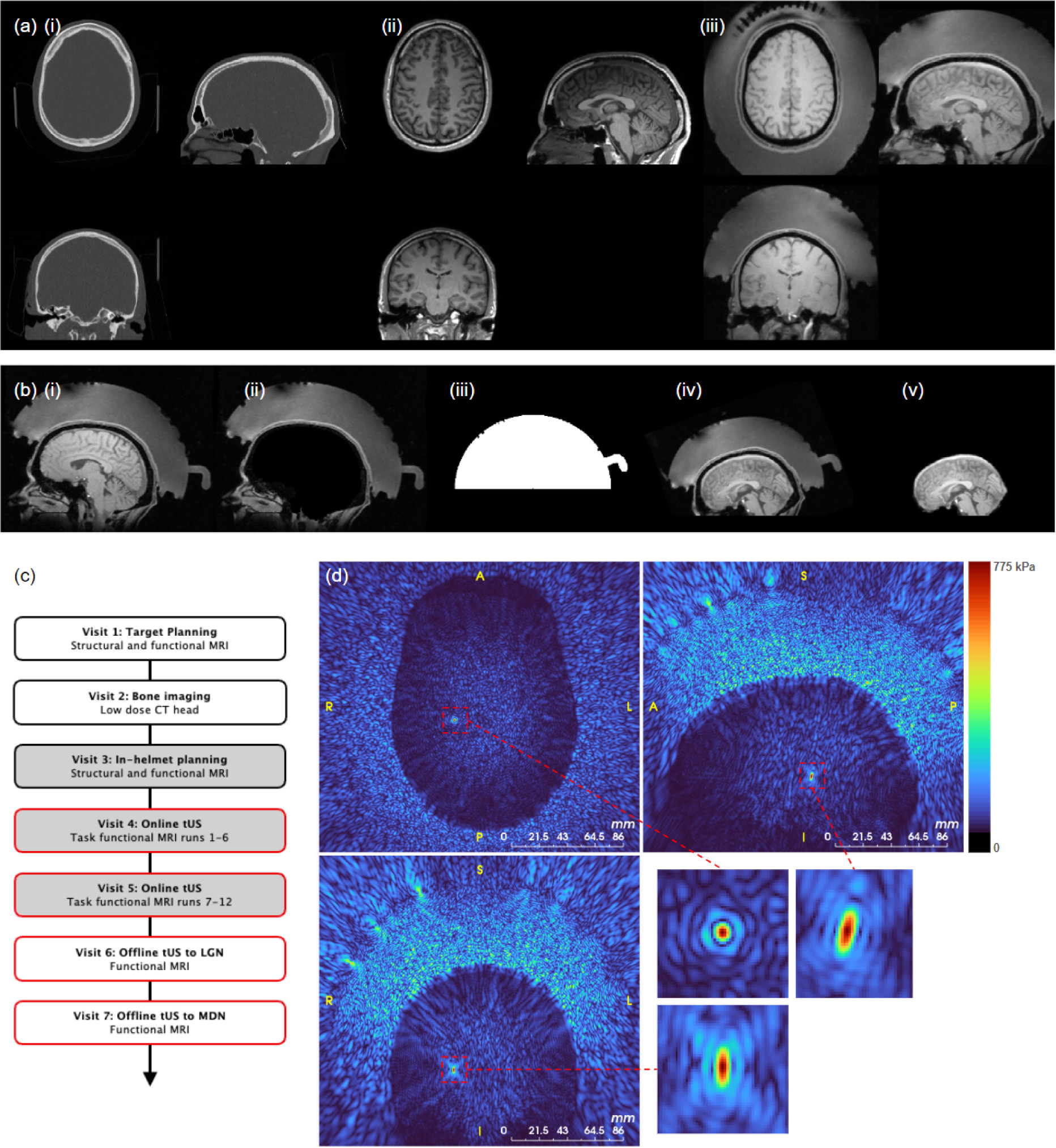
**(a)** Example planning images after registration (i) CT, (ii) planning T1 image obtained in the head coil, and (iii) positioning T1 image obtained in the helmet with no magnetisation pulse. **(b)** The positioning T1 image (i) is brain extracted (ii) and then registered with a reference image of the helmet (iii) to move the image to helmet space (iv). In the stimulation sessions, the brain is then registered with the brain from the positioning session (v), and the registration used to calculate any shift in the position of the focus. **(c)** Study visits. **(d)** Simulated acoustic pressure field for a target in the right LGN for one participant.

**Extended Data Figure 6:**
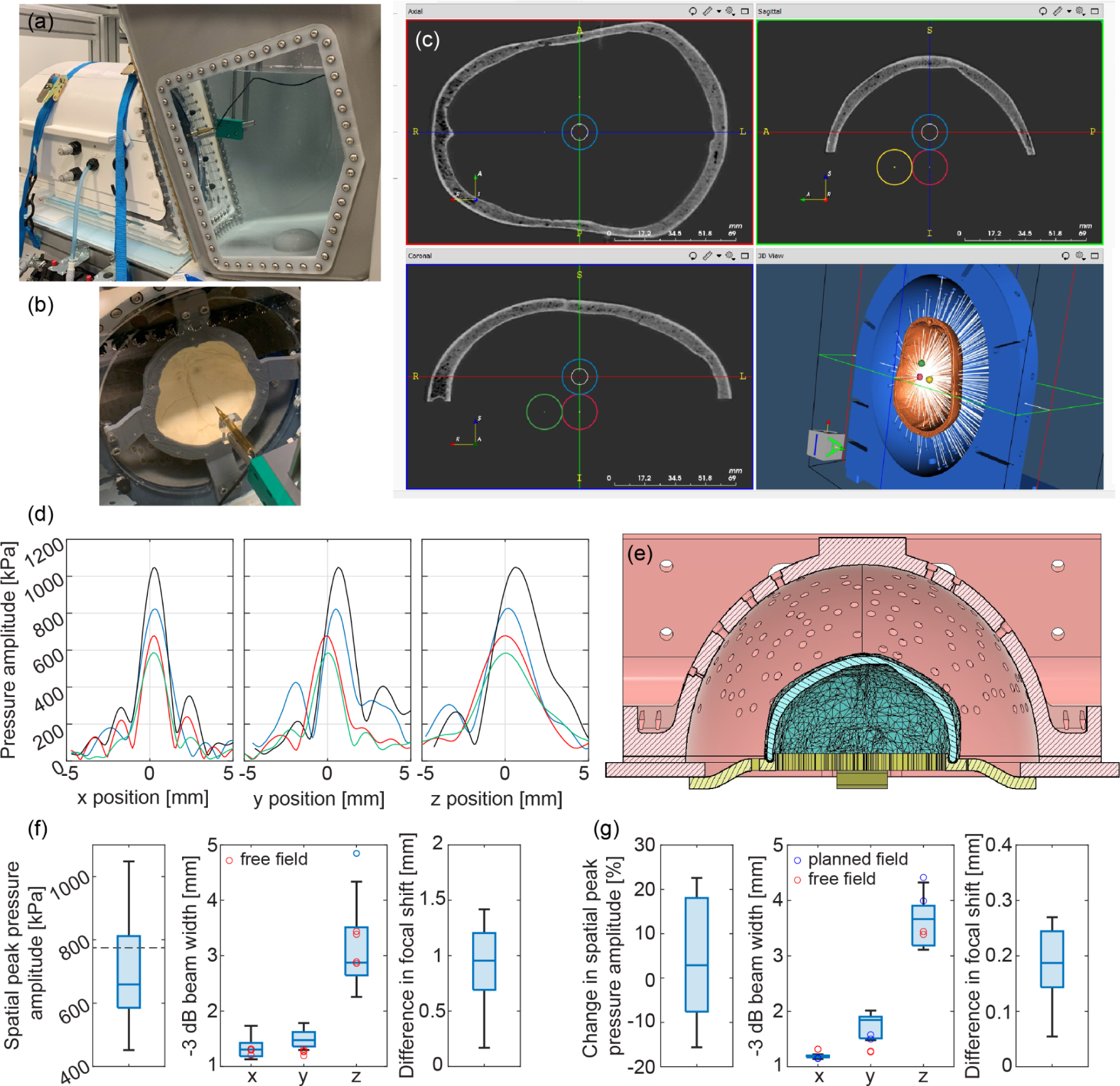
**(a)** Custom water tank field characterisation set-up. **(b)** Skull mounted in fixed position in array for skull validation measurements. **(c)** Screen-shot from k-Plan showing planned target positions inside an ex vivo human skull placed within the array. **(d)** Measured pressure amplitude profiles through the focus inside 4 skulls with planned position 0, 0, 0 and target pressure 750 kPa. **(e)** CAD drawing of the array housing with skull mount for registered geometry. The resulting stl file was used to register the position of the skull with the array elements for planning simulations. **(f)** Median and range (25th - 75th percentile) of spatial peak pressure amplitude, −3 dB beam width, and the position error (difference in focal shift) across 4 skulls with planned positions 0, 0, 0, 0, 0, 20 mm, 0, 20, 0 mm, 20, 0, 0 mm. The intended target pressure was 750 kPa (black dotted line). The red dots show the focal dimensions in free field demonstrating focusing capability is not affected by the presence of the skull after aberration correction. Difference in focal shift is calculated from the absolute difference between the intended shift (20 mm) and the euclidean distance to the measured focal position. **(g)** Accuracy of replanning showing change in the spatial peak pressure amplitude, the −3 dB beam width, and the position error (difference in focal shift) when replanning was performed to geometrically shift the focal position by 5 mm from the planned focal position. Red dots show the focal dimensions in free-field.

**Extended Data Figure 7:**
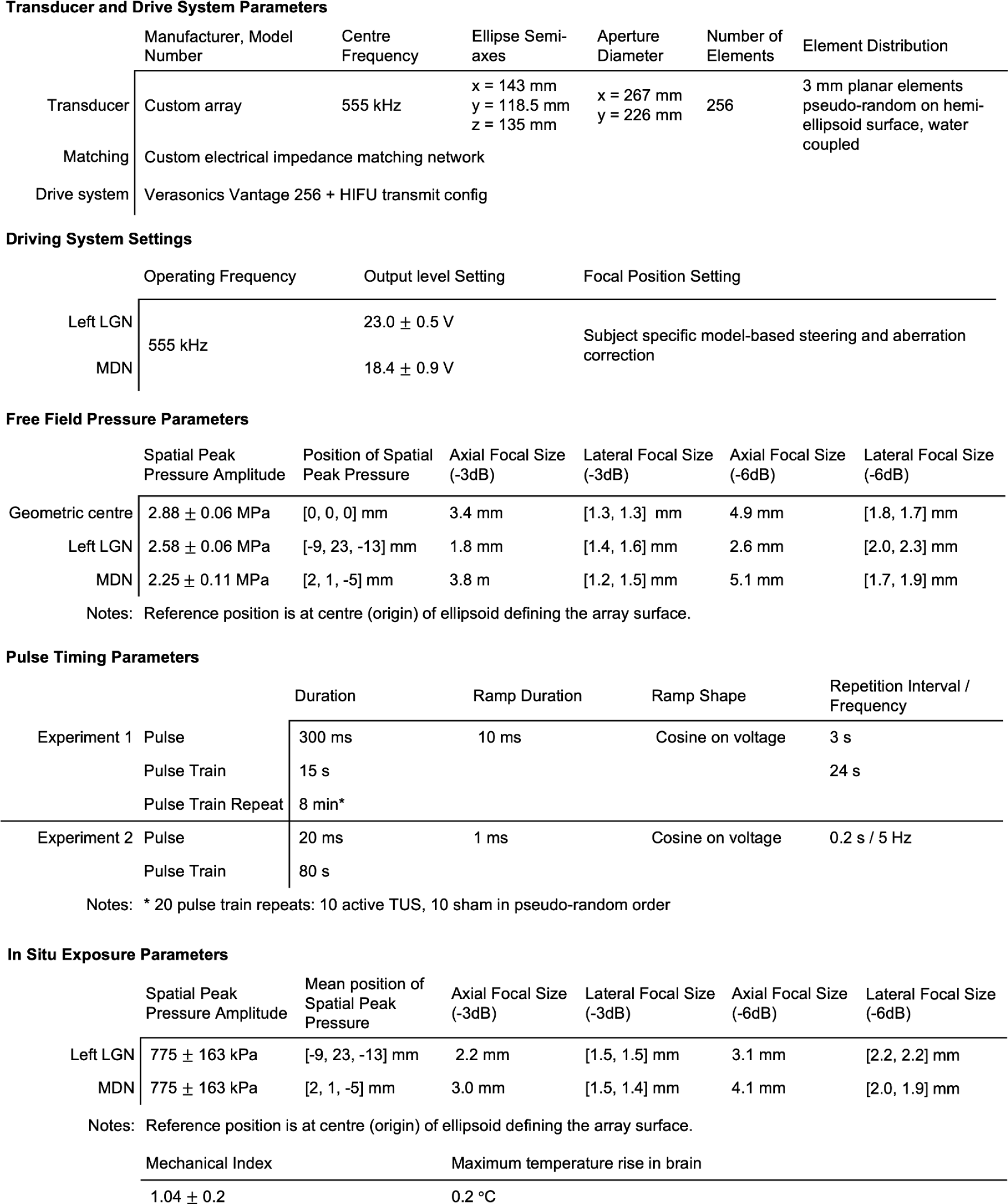
Summary of acoustic parameters for the study. Output level setting and free field spatial peak pressure amplitude is given as mean ± standard deviation across participants for each target. The position of spatial peak pressure for the LGN and MDN is the mean position across participants. The in situ estimates of spatial peak pressure were obtained from treatment planning simulations run in k-Plan and are given as planned pressure ± relative error in measured spatial peak pressure compared to simulation obtained from the skull validation measurements. This error was also used to define the uncertainty in the reported MI. The maximum temperature rise in the brain was the maximum value obtained from thermal simulations across all participants. The difference in lateral focal size between x and y dimensions and the differences in axial focal sizes between target positions arises in large part because the focal ellipsoid is not aligned with the helmet axes for all locations.

